# Emotion Dynamics as Hierarchical Bayesian Inference in Time

**DOI:** 10.1101/2021.11.30.470667

**Authors:** Gargi Majumdar, Fahd Yazin, Arpan Banerjee, Dipanjan Roy

## Abstract

What fundamental property of our environment would be most valuable and optimal in characterizing the emotional dynamics we experience in our daily life? Empirical work has shown that an accurate estimation of uncertainty is necessary for our optimal perception, learning, and decision-making. However, the role of this uncertainty in governing our affective dynamics remains unexplored. Using Bayesian encoding, decoding and computational modelling, we show that emotional experiences naturally arise due to ongoing uncertainty estimations in a hierarchical neural architecture. This hierarchical organization involves a number of prefrontal sub-regions, with the lateral orbitofrontal cortex having the highest representational complexity of uncertainty. Crucially, this representational complexity, was sensitive to temporal fluctuations in uncertainty and was predictive of participants’ predisposition to anxiety. Furthermore, the temporal dynamics of uncertainty revealed a distinct functional double dissociation within the OFC. Specifically, the medial OFC showed higher connectivity with the DMN, while the lateral OFC with that of the FPN in response to the evolving affect. Finally, we uncovered a temporally predictive code updating individual’s beliefs swiftly in the face of fluctuating uncertainty in the lateral OFC. A biologically relevant and computationally crucial parameter in theories of brain function, we extend uncertainty to be a defining component of complex emotions.

## Introduction

Imagine yourself in a close meeting with a friend only to know later that he had contracted COVID. The situation can lead to one of two possible outcomes – falling sick or remaining unaffected. Much of our real life emotional states are inescapably affected by shifting uncertainty of the future ^1^, and the drifting nature of this temporally, with changes in the environment characterizes our emotional experiences ^2^. However, this dynamicity is often overlooked while investigating affective experiences by transforming it into discrete emotional states ^3^. Although recent evidence suggests that emotions act on a continuous gradient^4,5^, most often it is reported as point estimates of specific emotion categories ^6^. The two underlying dimensions - valence and arousal - consistently identified to describe emotional experiences, are effective in defining disconnected feelings ^7^ but have been shown to be inefficient in characterising complex real-world emotions ^8,9^. To incorporate this complexity, varying combinations of context-driven dimensions like impulsivity, frustration, excitement have been used, which limits studies to describe occurrence of specific emotional states ^10–12^. Hence, these studies are not well suited to capture the fundamental properties of emotion. How then, can we formally characterize an emotional experience without excusing its real-world complexity and dynamicity?

A first principal approach would be to think from a fundamental property of the environment-experience interaction. All real-world social and emotional situations underlie a fluctuating uncertainty, with the outcome probabilities often incomputable as the causes are manifold and opaque ^13^. Our brain has to continuously generate predictions about the causes of these stimuli ^14–16^ and their future states and outcomes. Uncertainty defines the precision ^17^ with which a prediction can be made based on available information. Empirically, it is known to mediate the learning rate ^18,19^, synaptic plasticity ^20^, memory formation ^21,22^ and arbitrate between model-free and model-based systems ^23^. Thus, uncertainty acts as a biologically relevant and computationally crucial parameter for attention ^24,25^, inference ^17^, learning ^18,19,26^ and decision-making ^27–29^. Can it be the ubiquitous dimension in characterising our emotions incorporating both the complexity and the dynamicity?

Studies have shown that an accurate representation of uncertainty is essential for learning the statistical regularities in the model of the environment, and consequently, an improper representation can not only lead to learning deficits, but several emotional disorders like anxiety and depression ^30,31^. In major emotional disorders like anxiety ^32^, PTSD ^33^, panic disorders ^34^ and OCD ^35^, clinical studies have shown a pervasive role of lateral OFC. Interestingly, the OFC is a crucial region with distinct neuronal coding for expected uncertainty, as well as expected value of an outcome ^36–38^. Parallelly, a large number of prefrontal regions have been implicated in uncertainty processing in reward anticipation, uncertainty estimation ^37–39^ as well as emotion processing ^40,41^. Although these two empirical lines of work are examined separately, there is increasing evidence of neural overlap between the two ^41^. The involvement of other prefrontal areas like mPFC, dlPFC, ACC in processing uncertainty alongside OFC ^26,42^ with rich bidirectional connections with each other, entails the question of whether there is a distributed set of brain regions coordinating this processing ^40^, with underlying local and global computations for the resolution of uncertainty. Furthermore, it is currently unknown whether the role of OFC in emotions is to represent uncertainty about future states.

We formally addressed the role of uncertainty in emotional states and its underlying neural correlates by using a well-studied naturalistic movie with socially charged and uncertain outcomes. This uncertainty, reflected by the variance between its outcomes, fluctuated dynamically and was quantified by an independent group of human raters. Using hierarchical Bayesian modelling, we found that uncertainty can explain and influence emotional intensity significantly more than valence. The widespread cortical activations due to uncertainty as compared to valence also substantiated this. Crucially we uncovered a specific role for the lateral OFC, which was parametrically modulated by uncertainty, and in response, showed more complex neural representations accordingly. This neural response towards uncertain events by participants could significantly predict their baseline anxiety scores. The temporal dynamics of uncertainty processing demonstrated a functional double dissociation between the lateral and medial OFC. The former integrated uncertainty dynamically with the Fronto-Parietal system, while the latter showed temporal functional integration with the Default Mode Network. Generative modelling of neural and behavioural data revealed that uncertainty (and arousal) can naturally arise from estimations of volatility in valence, and temporal updates to this were seen in the ongoing neural states in the lOFC. This expands our understanding of the functional role of OFC in emotions in humans and necessitates the incorporation of uncertainty as a fundamental dimension to real-world emotions.

## Results

### Behavioural ratings

#### Quantifying uncertainty, valence, and arousal dynamically

For our study, we operationalised Uncertainty as outcome uncertainty ^13^ where larger variance between the values of the possible outcomes (e.g. shoot vs no shoot) increases the perceived uncertainty ^30^. This captures the dynamicity in temporally extended stimulus like ours, with outcomes unfolding over a period.

We characterised the emotion dynamics by measuring the continuous rating of the movie for Arousal, Valence ^43^ and Uncertainty from 3 independent groups of participants (n = 60, 20 participants per group). To assess the reliability of their responses we performed an ISC ^44^ for each group (Fig. 1a) and excluded participants with r < 0.2. A strong positive ISC for each group (Mean ISC: r = 0.62, SD = 0.14 (Arousal); r = 0.73, SD = 0.15 (Valence); r = 0.57, SD = 0.19 (Uncertainty)) confirmed a consistent shared response for all behavioural measures across participants.

**Figure 1:**
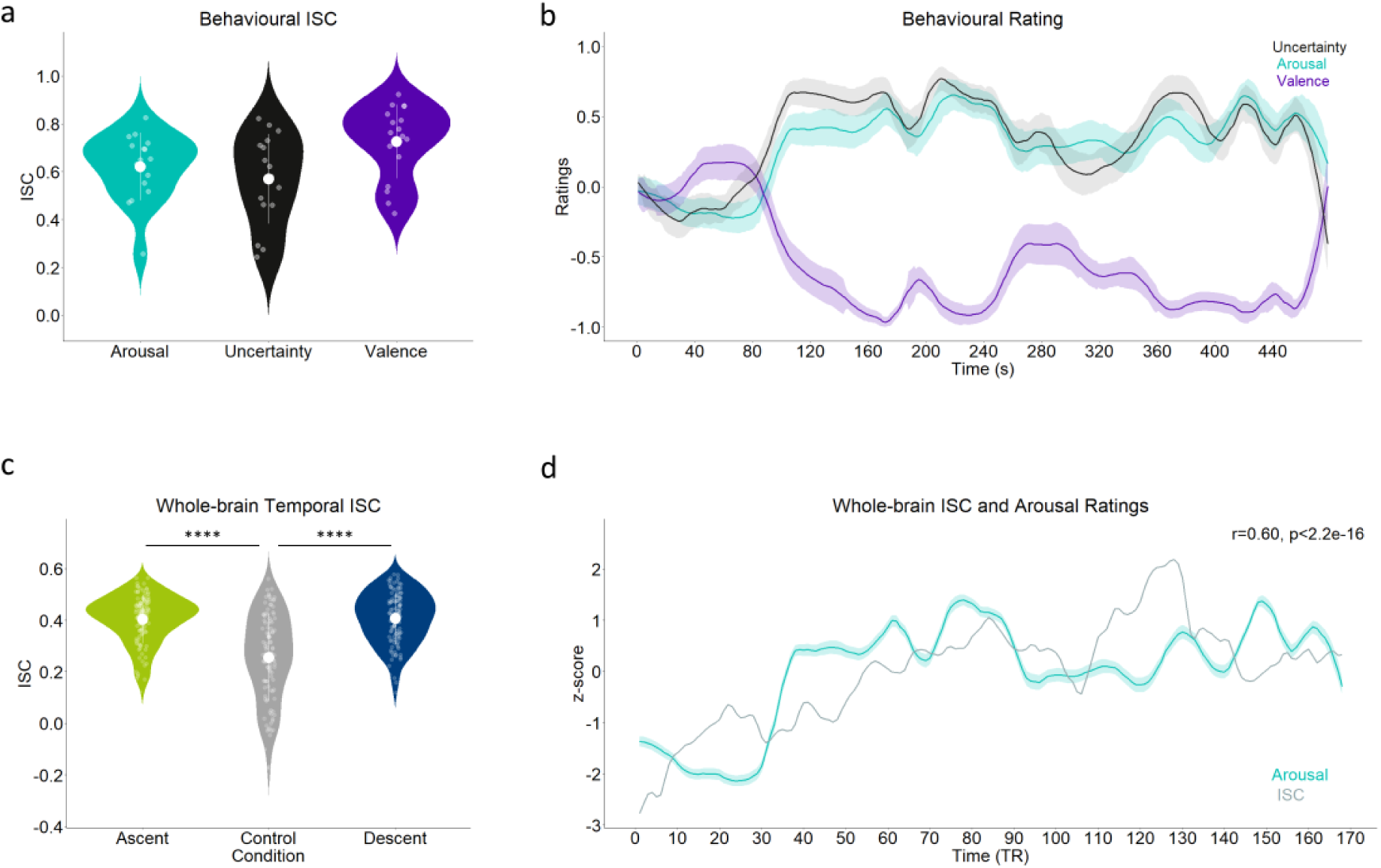
Neural engagement reflects behavioural responses. (**a**) Violin plots show the mean behavioural ISC for the whole movie for 3 measures – Arousal (r = 0.62, n = 17), Uncertainty (r = 0.57, n = 16) and Valence (r = 0.73, n = 17). White circular markers denote the mean ISC for each, with white bars as the SD. Scattered dots represent individual subject’s ISC values. (**b**) Timeseries of participants’ continuous affect ratings across the movie showing strong positive correlation between Arousal-Uncertainty (r = 0.86, p < 0.0001) and strong negative correlation between Arousal-Valence (r = -0.95, p < 0.0001). Shaded regions represent the SE at each second. 5 Ascent, 5 Descent and 1 Control segment were isolated from the Arousal ratings. The BOLD signals were then concatenated to obtain one time series for each condition from the fMRI participants. (**c**) Violin plots show the whole-brain ISC of fMRI participants (n = 111) for 3 conditions – white circular dot marks the average ISC for Ascent (r = 0.40), Control (r = 0.26) and Descent (r = 0.40) indicating significant difference between both Ascent and Descent with Control. Scattered dots represent individual subject’s ISC values. (**d**) Average whole-brain ISC (z scored) computed over 21TR sliding window (with a step-size of 1TR) shows strong positive correlation with the smoothed, downsampled and z-scored Arousal ratings of behavioural participants (r = 0.60)

#### Defining emotionally dynamic conditions

The behavioural ratings of Arousal revealed that the movie elicited strong and temporally dynamic emotional arousal (Mean = 0.35) (Fig. 1b), ranging from (−0.30 to 0.81). Arousal time course was taken as a proxy of emotional experience to identify emotionally dynamic conditions for our main analyses. We specified an Ascent phase (signifying the rising intensity in the anticipatory phase), a Descent phase (signifying the falling intensity with the release of uncertainty) and a Control phase. A total of 5 Ascents (Mean duration = 21 sec, SD = 6.3), 5 Descents (Mean duration = 16 sec, SD = 5) and one Control (Duration = 25 sec) segment were identified for successive analyses (See methods).

#### Relationship between the behavioural measures

Next, we tested the relationships between the behavioural measures and found a strong positive correlation between Arousal and Uncertainty (r = 0.86, p = 4.31e-141) (Fig. 1b). Furthermore, a strong anti-correlation (r = -0.95, p = 1.65e-248) between Valence and Arousal was also observed (Fig. 1b), consistent with previous literature ^43^. Although the ratings were not given by the same individuals from the CamCan dataset, 86% of the included participants had 50% or above intersubject agreement suggesting robust responses.

### Neural engagement reflects behavioural engagement

Since the behavioural responses were collected from independent groups of participants, we wanted to see whether the behavioural experience is reflected by the neural engagement of the CamCan population. For this we performed a whole-brain spatial ISC (Whole brain parcellation by Power Atlas, 264 ROIs) ^45^, for the fMRI participants (n=111) and correlated that with the smoothed and downsampled Arousal ratings (Fig. 1d), as high arousal shows high neural ISC ^43^. We indeed found a significant positive correlation (r = 0.60, p = 4.17e-18) supporting our hypothesis that neural engagement increased and decreased with varying Arousal in time, consistent with previous findings in literature ^43^.

### Validating the emotional saliency of the Conditions

Thereafter, we confirmed the emotional saliency of our conditions by performing a whole-brain temporal ISC for each condition (Ascent, Descent and Control) (Fig. 1c). We found both Ascent and Descent to have significantly higher ISC than Control (Ascent: r = 0.40, SD = 0.09; Descent: r = 0.40, SD = 0.10; Control: r = 0.26, SD = 0.17; Ascent>Control: t = 9.14, p = 3.49e-15, Cohen’s d = 1.06; Descent>Control: t = 9.14, p = 3.49e-15, Cohen’s d = 1.08) proving that they are emotionally charged scenes. Furthermore, we extended this analysis to each scene to resolve any underlying bias of the scenes’ duration or content (Supplementary Fig. 1). Each Ascent and Descent showed the same pattern when contrasted with Control, validating the stimulus properties regarding emotional dynamics and the synchronicity between observed neural signal and rated affective engagement.

### Arbitrating between the effect of Uncertainty and Valence

A large body of evidence suggests our brain employs Bayesian inferences for a wide range of cognitive processes ^14,20,26, 46–48^, with uncertainty acting as the driving parameter. In the same vein, one can assume that affective dynamics would be no exception. We hypothesised that in situations of varying emotional outcomes (e.g., shoot or no shoot), the outcome uncertainty would be playing a more significant role in the emotional dynamics than valence (i.e., negative or positive).

To delineate the effects of Uncertainty and Valence on emotion dynamics, we constructed a Bayesian Hierarchical Regression model ^49^ specifying the effect of Uncertainty and Valence as their interaction with Arousal (Fig. 2a). This model accounted for subject variability, content variability, ROI-level variability, and subject sensitivity to Uncertainty/Valence/Arousal (See methods). Crucially, unlike point estimates, this model gives a full posterior distribution enabling us to do robust estimations using a statistically conservative approach (See methods for model specifications including choice of priors). Model validation using posterior predictive check showed the model was a good fit for the data (Supplementary Fig. 3a, b).

**Figure 2:**
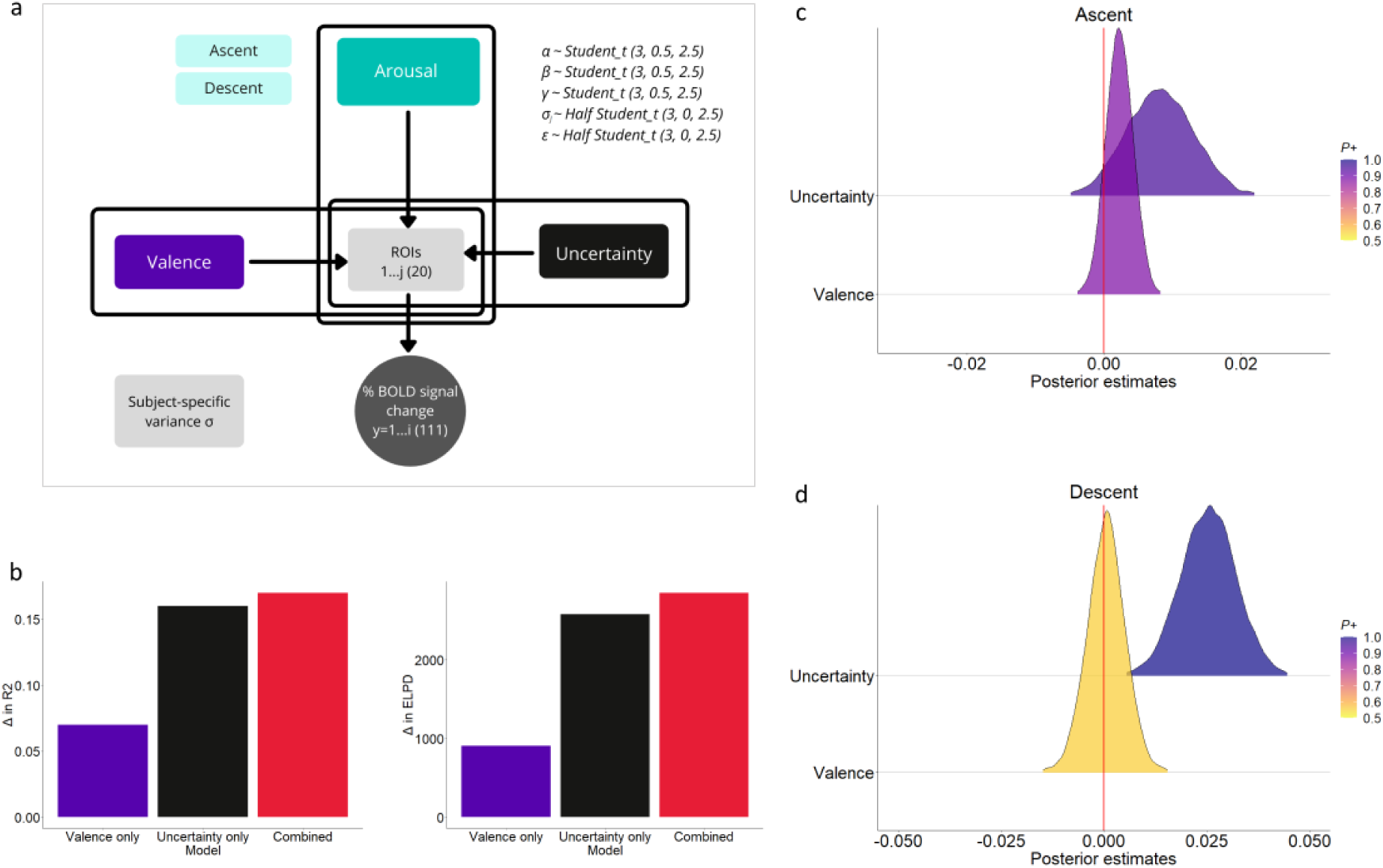
Arousal related neural activations are better explained by Uncertainty than Valence. **(a)** Graphical illustration represents the architecture of the Hierarchical Bayesian model. % BOLD was modelled to be drawn from a Student t-distribution and varied across participants and ROIs. Main effects of interest were the interaction terms of Arousal (as categorical predictor with two levels) with each of Uncertainty and Valence (as continuous predictors), which were drawn from a noninformative Student t prior. The slopes of these varied across each ROI to extract ROI-level inference. (**b**) Changes in Bayesian-R^2^ (left) and expected log predictive density (right) for the Valence-only model (purple), Uncertainty-only model (black) and combined (red), compared to the null model. These metrics quantify the quality of model fit and predictive capability of the model, respectively. (**c**) Posterior density estimates for Ascent showing the effect of Uncertainty (P = 0.95) has more evidence than Valence (P = 0.88). (**d**) Posterior density estimates for Descent showing the effect of Uncertainty (P = 0.99) has more evidence than Valence (P = 0.56).

First, we ran a null model where the % BOLD of the ROIs are explained only by Ascent and Descent phases. Thereafter, we ran three more models - Uncertainty-only model (UncM), Valence-only model (ValM) and a final model including both Uncertainty and Valence (Combined), to measure each parameter’s unique contribution in explaining the data. Model performance was assessed by their R^2^ and expected log predictive density (elpd) values (relative to the null model) (Fig. 2b). The resulting gain in R^2^ (ΔR^2^) (Fig. 2b<u>, left</u>) or elpd (Δelpd) (Fig. 2b<u>, right</u>) relative to the null model provides an upper bound of unique information contributed by Uncertainty or Valence in explaining the data, in addition to Arousal. Results showed that UncM possessed more than twice the explanatory power (ΔR^2^ = 0.16, Δelpd = 2583.4) than ValM (ΔR^2^ = 0.07, Δelpd = 907.4) in both the measures of model comparison, which were only modestly below the Combined model (ΔR^2^ = 0.17, Δelpd = 2854.4). This strongly suggests that uncertainty plays a significant role in emotion dynamics.

We selected the Combined model as our final model to inspect the main interaction effects of Uncertainty and Valence for Ascent and Descent. Model output showed that the posterior evidence (P) of the main effect of Uncertainty was significantly more than Valence in Ascent (Uncertainty β_mean_ = 0.008, 95% HDI = [-0.001, 0.018], P = 0.95, Valence β_mean_ = 0.002, 95% HDI = [ -0.001, 0.006] , P = 0.88) (Fig. 2c) as well in Descent (Uncertainty β_mean_ = 0.025, HDI = [0.012, 0.039], P = 0.99, Valence β_mean_ = 0.0005, 95% HDI = [-0.009, 0.009], P = 0.56) (Fig. 2d). An additional one-sided hypothesis testing (analogous to GLM ‘contrasts’) demonstrated the effect of Uncertainty to be marginally more than Valence in Ascent (P = 0.9, BF = 9.31) and significantly more in Descent (P = 0.99, BF = 1999).

It should be noted that the change in valence across the scenes for Ascent were not sufficient for the model to show a significant effect. This crucially reflects a lower sensitivity of valence in affecting arousal, implying the necessity of a higher dynamic range of valence to elicit significant effect, which makes it less reliable as a predictor for nuanced changes ^8^, compared to uncertainty. Moreover, Descent, though characterised by decreasing Uncertainty, showed more evidence for Uncertainty than Valence, validating those small changes in uncertainty can contribute to a significant effect than the corresponding increase in valence, even for falling intensities of emotion. Taken together, this clearly outlines the substantive role of uncertainty in explaining neural activations due to emotional experiences.

### Uncertainty parametrically modulates more regions than valence

Next, we sought to estimate and quantify the effect of Uncertainty and Valence in modulating the % BOLD of the ROIs (signifying ROI-level effect or “group-level effects” in the mixed-model parlance) during Ascent and Descent from our Bayesian model (Fig. 3). This approach allowed us to quantitively model the bidirectional effect of the parameters on the ROIs by computing the posterior evidence for each ROI.

**Figure 3:**
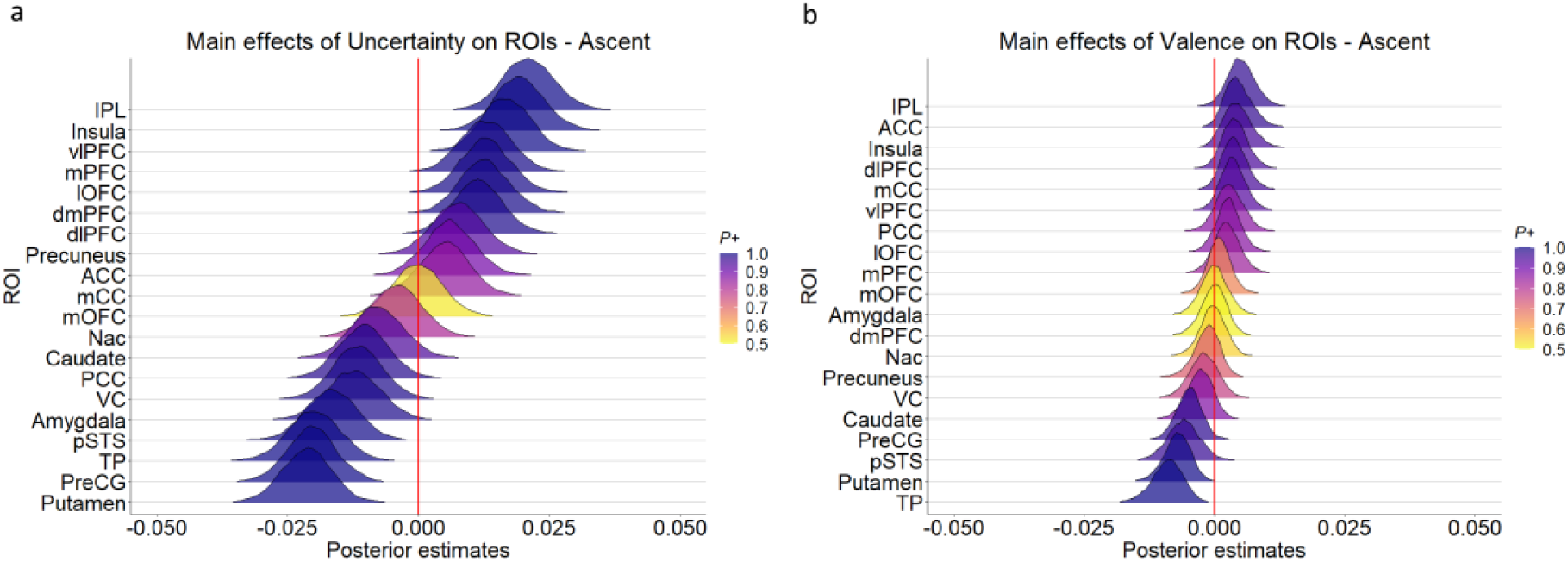
Uncertainty parametrically modulates more regions compared to Valence. (**a**) Posterior density plots show the effect of Uncertainty on all ROIs during the Ascent phase. Areas showing evidence of increasing activity with Uncertainty include IPL, anterior Insula, vlPFC, mPFC, lOFC, dmPFC and dlPFC. (**b**) Posterior density plots showing the effect of Valence on all ROIs during the Ascent phase. Areas showing evidence of increasing activity with Valence include ACC, IPL and anterior Insula.

We focused our results on the Ascent phase as it represents heightened intensity of emotional experience. Uncertainty showed a widespread effect on both cortical and subcortical regions compared to Valence revealing a distinct asymmetry of activity pattern. Most prefrontal ROIs including lOFC (β_mean_ = 0.013, 95% HDI = [0.003, 0.022],P = 0.99), dmPFC (β_mean_ =0.013, HDI = [0.003, 0.022], P = 0.99), dlPFC (β_mean_ = 0.011, HDI = [0.002, 0.021], P = 0.99), vlPFC (β_mean_ = 0.017, HDI = [0.008, 0.026], P = 0.99), mPFC (β_mean_ = 0.013, HDI = [0.003, 0.022], P = 0.99), ant INS (β_mean_ = 0.019, HDI = [0.009, 0.029], P = 0.99) along with IPL (β_mean_ = 0.021, HDI = [0.011, 0.031], P = 0.99) exhibited strong evidence for increased activation with increased Uncertainty in Ascent (Fig. 3a). In contrast, only ACC (β_mean_ = 0.005, HDI = [0.000, 0.009], P = 0.98), ant INS (β_mean_ = 0.004, HDI = [-0.0006, 0.01], P = 0.95) and IPL (β_mean_ = 0.005, HDI = [0.000, 0.01], P = 0.98) showed evidence for increased activation with increasing Valence (Fig. 3b).

The model also revealed evidence for increased activation in Amy (β_mean_ = -0.012, HDI = [- 0.022, -0.002], P = 0.99), pSTS (β_mean_ = -0.016, HDI = [-0.026, -0.006], P = 0.99), TP (β_mean_ = - 0.019, HDI = [-0.03, -0.01], P = 0.99), PreCG (β_mean_ = -0.02, HDI = [-0.03, -0.01], P = 0.99), Put (β_mean_ = 0.021, HDI = [-0.03, -0.012], P = 0.99), PCC (β_mean_ = -0.010, HDI = [-0.02, -0.001], P = 0.99) and VC (β_mean_ = -0.011, HDI = [-0.021, -0.002], P = 0.99) with decreasing Uncertainty (Fig. 3a). For Valence, TP (β_mean_ = -0.009, HDI = [-0.015, -0.003], P = 0.99), Put (β_mean_ = -0.007, HDI = [-0.012, -0.002], P = 0.99), pSTS (β_mean_ = -0.006, HDI = [-0.012, 0.0002], P = 0.97) and PreCG (β_mean_ = -0.005, HDI = [-0.009, -0.0002], P = 0.97) also showed evidence for increased activation with decreasing Valence (Fig. 3b).

Our findings on the effect of Uncertainty and Valence in modulating the neural activity during Ascent revealed a distinct asymmetry with a widespread activity for Uncertainty compared to Valence. Even the same regions showed greater modulation by Uncertainty than by Valence. Furthermore, it outlines a crucial role of the prefrontal regions in mediating uncertainty during increased arousal. The extensive effect of Uncertainty on cortical and subcortical ROIs was also observed during Descent (Supplementary Fig. 4a).

### Temporal dynamics track temporal Uncertainty

Specifying Ascent and Descent for our Bayesian model makes it a supervised approach to discern the effect of the parameters. To address this bias that may constrain our inference, we employed a spatiotemporal PCA ^50,51^ to the BOLD time series of the ROIs to delineate the effect of Ascent and Descent in a data-driven way from the resulting components. First, we performed a spatial PCA on individual subject’s multivariate, scene-averaged BOLD time series to obtain the weights associated with each ROI. We selected the first 2 PCs of the spatial PCA, which accounted for 71% of the total variance, for our further analyses (Fig. 4e).

**Figure 4:**
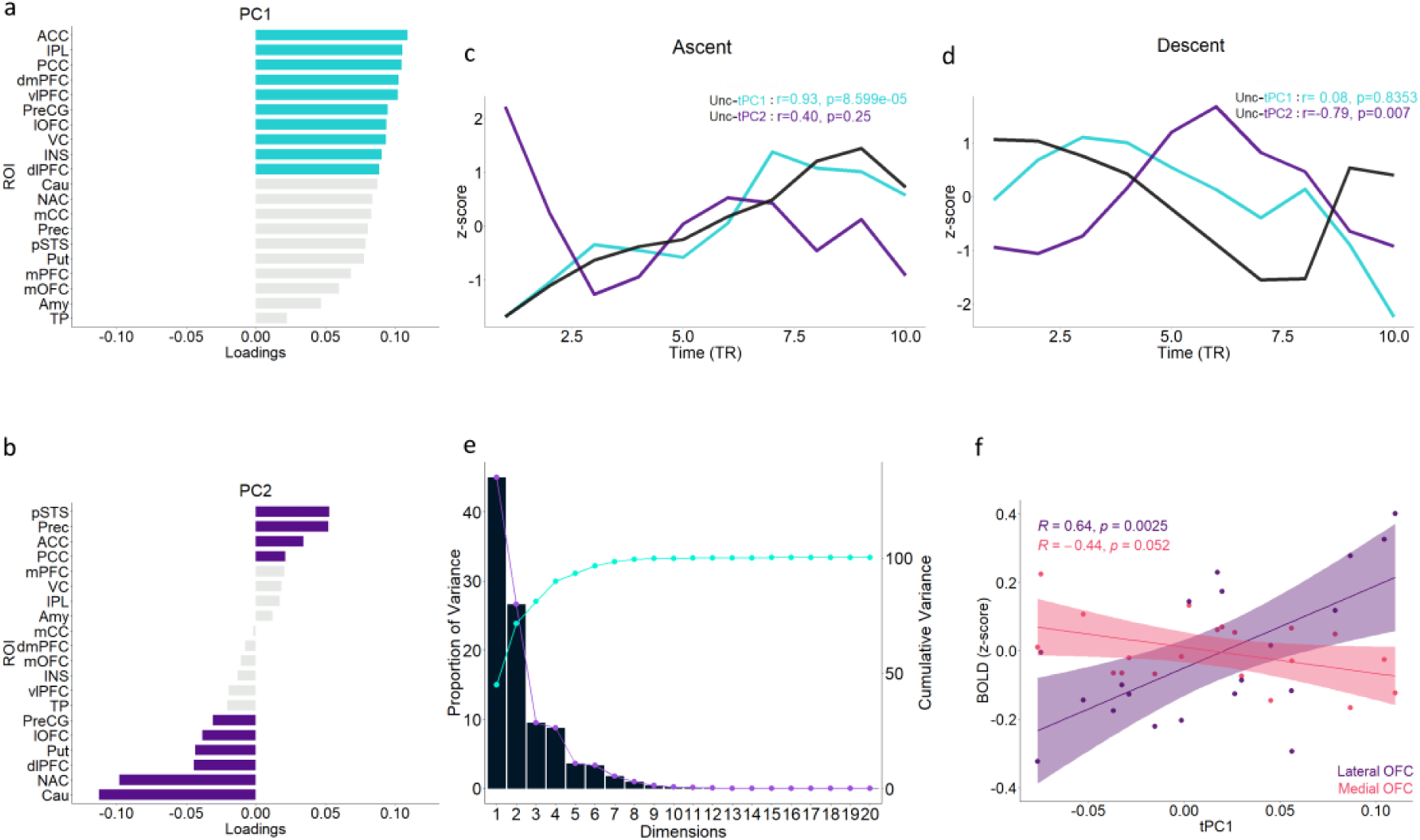
Low dimensional neural signal tracks temporal Uncertainty. (**a**) First principal component (PC1) showing the loadings of all the ROIs. The top ten contributing ROIs (shaded cyan) include all prefrontal regions, except mPFC and mOFC. (**b**) PC2 showing the loadings of all the ROIs, with top ten contributors shaded purple. (**c**) The correlation of averaged Uncertainty ratings of Ascent with first and second temporal principal components (tPC1 and tPC2) reflecting significant correlations with tPC1 but not tPC2. (**d**) In Descent, tPC2 but not tPC1 shows a significant correlation with mean Uncertainty ratings. (**e**) Scree plot: Line plot (purple) representing the percentage of variance explained by the PCs; Line plot (cyan) showing the cumulative variance across PCs; Bar plots depicting the percentage of total variance explained by the PCs. (**f**) Pearson correlation of z-scored BOLD signal of lOFC and mOFC with tPC1.

Next, to assess the temporal evolution of these two components, we performed a temporal PCA by computing the dot product of the PC weight vectors with the ROI BOLD time series. This yielded corresponding tPCs, which were then averaged across the subjects to obtain the group-level tPCs. tPC1, which explained 45% of signal variance, showed a very strong correlation with the Ascent Uncertainty (r = 0.93, p = 0.00009) but not with Descent Uncertainty (r = 0.075, p = 0.836) (Fig. 4c). However, tPC2 did not show any significant correlation with Ascent Uncertainty (r = -0.399, p = 0.253), while displaying a strong negative correlation with the Descent Uncertainty (r = -0.785, p = 0.007) (Fig. 4d). The negative direction of this component correlation suggests that there is indeed an increase in (average) neural activity in Descent (as opposed to the downward direction of Uncertainty). Thus, identifying the spatiotemporal PCs provided evidence for lower dimensional temporal dynamics of Ascent and Descent reflected by tPC1 and tPC2, respectively.

The PC1 loadings (Fig. 4a) revealed a large contribution of prefrontal areas besides IPL, VC and PCC. Although a caveat of caution is advised for the correlation result (because of the lesser number of time points), the fact that the majority of the areas (9/10) contributing to PC1 were also modulated by uncertainty is a strong corroboration of the Bayesian model output.

Given the extensive evidence of the role of OFC in future state prediction ^52^, encoding latent state ^53^ and processing uncertainty ^13,54–56^ in other cognitive domains, we next hypothesised that OFC might be tracking the uncertainty in affective experiences. However, converging evidence from the Bayesian model and the Spatiotemporal PCA showed evidence for lateral but not medial OFC to be affected by uncertainty. To examine whether this pattern is reflected temporally as well, we correlated the lOFC and mOFC BOLD signals with tPC1 and found a significant positive correlation with lOFC (r = 0.64, p = 0.003) but not with mOFC (r = -0.44, p = 0.05) (Fig. 4f).

Taken together, this enabled us to distinguish between the conditions, identify an affectively relevant neural signal showing large contribution by the PFC (supporting the Bayesian model’s inference) and helped us uncover a distinct temporal signature between lOFC and mOFC in processing uncertainty in emotional experiences.

### A functional double dissociation between lOFC and mOFC in processing uncertainty

To further investigate the functional differences between the lOFC and mOFC as evidenced by the tPCA, we performed an ISFC of lOFC and mOFC with other ROIs over a sliding window of 21 TR to identify stimulus-locked network correlation patterns shared across participants. We categorised our ROIs into three networks – DMN (mOFC, PCC, Prec), FPN (dlPFC, vlPFC, IPL) and Limbic (Amy, Put, Nac, Cau) since these areas showed an opposing pattern in our model output as well as serve as core nodes of canonical functional networks. Thereafter, we correlated the ISFC correlation coefficients with the Uncertainty and Valence time series and applied a multi-dimensional scaling on the resultant values (See methods). This enabled us to estimate the correlation strength between the nodes and importantly examine the nodes that cluster together given a parameter.

Figure 5a (left) illustrates that lOFC, but not mOFC tend to cluster more with the nodes of FPN showing increased ISFC with Uncertainty (lOFC-dlPFC: r = 0.46, p = 4.6e-10, lOFC-vlPFC: r = 0.65, p = 2.2e-16, lOFC-IPL: r = 0.54, p = 4.4e-14). A completely opposite pattern was observed for mOFC which clustered strongly with the DMN nodes (mOFC-mPFC: r = 0.78, p = 2.2e-16, mOFC-PCC: r = 0.44, p = 2.7e-9, mPFC-Prec: r = 0.51, p = 2.8e-12) (Fig. 5c<u>, left</u>). Figure 5b and Figure 5d show ISFC timeseries of lOFC and mOFC with vlPFC and mPFC respectively with the Uncertainty. Both lOFC and mOFC showed widespread correlation with the subcortical ROIs with no distinct pattern (Supplementary Fig. 5). For all networks the ISFC-Valence correlation values showed an opposite pattern supporting the anticorrelation observed between Uncertainty and Valence in the behavioural analysis. These results suggest a double dissociation of OFC with possible role of lOFC in encoding the external uncertainty in concert with similarly oriented ROIs. Conversely, the involvement of mOFC with self-referential areas of the DMN might be reflecting its role in endogenous processing of uncertainty ^57^.

**Figure 5:**
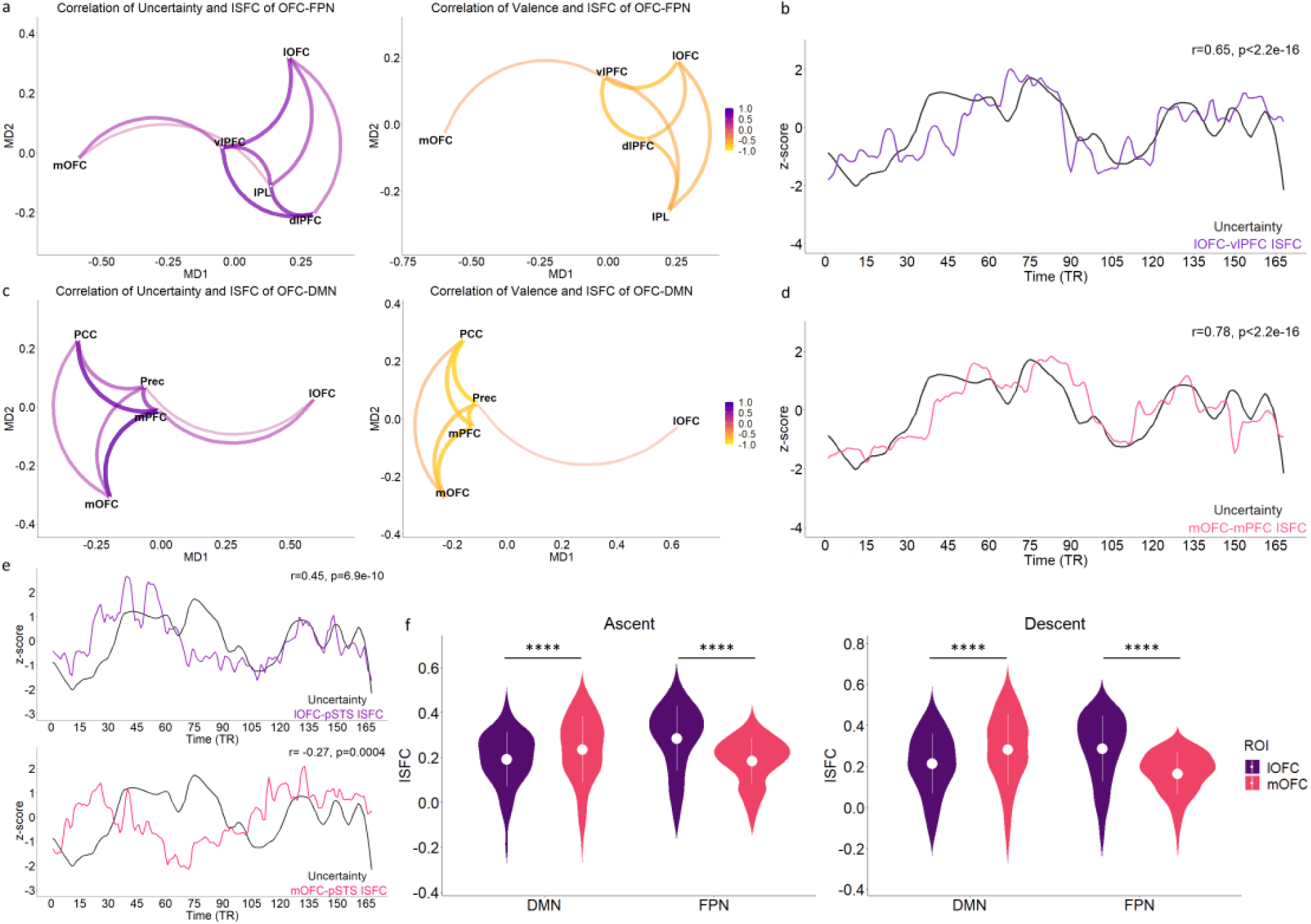
A functional double dissociation between lOFC and mOFC. (**a**) & (**c**) depicts multidimensional scaling of correlation coefficients obtained by correlating sliding window ISFC time series of an ROI pair with averaged Uncertainty ratings. The line joining two nodes (or ROIs) represents the correlation strength between their ISFC and Uncertainty. Nodes clustered together show similar correlations. (**a**) outlines clustering of lOFC but not mOFC with the FPN regions, whereas (**c**) reflects the opposite pattern in the case of DMN nodes. A similar pattern is seen when the ISFCs are correlated with Valence for both the networks (right). (**b**) & (**d**) shows ISFC between ROIs having highest correlation with Uncertainty for each network. Mean sliding window ISFC (z-scored) between (**b**) lOFC-vlPFC (r = 0.65) and (**d**) mOFC and mPFC (r = 0.78) showing significant correlation with average ratings of Uncertainty (z-scored and downsampled) (**e**) Differential processing of uncertainty by lOFC and mOFC (external versus internal respectively) validated by correlating their ISFC time courses with pSTS and Uncertainty. lOFC-pSTS showed a significant positive correlation (r= 0.45) with Uncertainty (top) while that of mOFC-pSTS revealed a significant anticorrelation (r = -0.27) (bottom). (**f**) Violin plots depicting the temporally averaged ISFC of lOFC and mOFC as the seed with DMN and FPN separately for Ascent (left) and Descent (right) conditions, reflecting the dynamic connectivity and MDS results.

To validate this, we extended this analysis to assess their connectivity with pSTS – a region, majorly associated with external dynamic stimuli processing. We found that lOFC had significant positive correlation with pSTS (r = 0.45, p = 6.86e-10) (Fig. 5e<u>, top</u>) while mOFC showed an opposite trend (r = -0.27, p = 0.0004) (Fig. 5e<u>, bottom</u>).

The above results clearly demonstrate a functional differentiation of lOFC and mOFC in processing uncertainty across time. Next, we assessed whether this differentiation is reflected when we isolate the Ascent and Descent. For that, we performed ISFC separately on the Ascent and Descent timepoints to examine the correlation pattern of lOFC and mOFC with DMN and FPN. This enabled us to precisely tease out the time points of Ascent and Descent (as there is no overlapping time-window) and assess the correlation. We found that ISFC between lOFC and FPN was significantly higher than that of mOFC and FPN for Ascent (lOFC-FPN: Mean r = 0.29, SD = 0.14; mOFC-FPN: Mean r = 0.18, SD = 0.24; t = 19.12, p = 9.18e-37, Cohen’s d = 0.57) (Fig. 5f<u>, left</u>). The reverse pattern was observed for DMN (lOFC-DMN: Mean r = 0.19, SD = 0.12; mOFC-DMN: Mean r = 0.24, SD = 0.15; t = 8.68, p = 3.98e-14, Cohen’s d = 0.29) (Fig. 5f<u>, left</u>). Crucially the same pattern was observed for Descent as well wherein FPN had significantly higher ISFC with lOFC than mOFC (lOFC-FPN: Mean r = 0.28, SD = 0.16; mOFC-FPN: Mean r = 0.16, SD = 0.10; t = 18.56, p = 1.09e-35, Cohen’s d = 0.51) (Fig. 5f<u>, right</u>) and DMN exhibited significantly higher ISFC with mOFC than lOFC (lOFC-DMN: Mean r = 0.21, SD = 0.15; mOFC-DMN: Mean r = 0.28, SD = 0.17; t = 12.33, p = 1.77e-22, Cohen’s d = 0.4) (Fig. 5f<u>, right</u>).

In summary, we posit a functional dissociation within the OFC in processing uncertainty in affective experiences, where lOFC is modulated by externally mediated cues about uncertainty and mOFC is mainly involved with possibly processing endogenous cues.

### Hierarchical organisation in the complexity of representation of uncertainty

#### Temporal autocorrelation

The results thus far inform us about the fundamental role of uncertainty driving emotional arousal, primarily mediated by the prefrontal regions with a functional double dissociation of lOFC and mOFC reflecting the exogenous and endogenous processing of affective uncertainty. We next carried out a series of analyses aimed at understanding how the representational complexity of uncertainty differs between OFC and the rest of the prefrontal areas.

First, we hypothesised that if there exists a hierarchy of complexity of representation among these regions, it would be reflected in their rate of information integration across the movie. Since the level of uncertainty dynamically varies with time in this movie, areas integrating information over a shorter span would fail to capture this complexity and represent low-level features of the stimulus. To quantify this rate of temporal integration, we estimated the temporal autocorrelations of each ROI’s voxel-wise time series for the whole movie. The decay of this signal with increasing lags (up to 50 TRs) evidently showed a hierarchy of timescales within the prefrontal cortex. We then fit an exponential function to the median of these estimates for each ROI using a robust estimator and calculated the number of time points before the function decayed to an arbitrary threshold of 0.2 (Fig. 6a). We found that medial and lateral OFC had the longest decay (mOFC = 28 TR, lOFC = 25 TR) compared to other prefrontal areas (mPFC = 21 TR, dmPFC = 16 TR, ACC = 14 TR, dlPFC = 12 TR, vlPFC = 12 TR). Our findings thus demonstrate that the prefrontal areas exhibit an intrinsic hierarchy across time, with the OFC occupying the highest level while processing naturalistic affective experiences.

**Figure 6:**
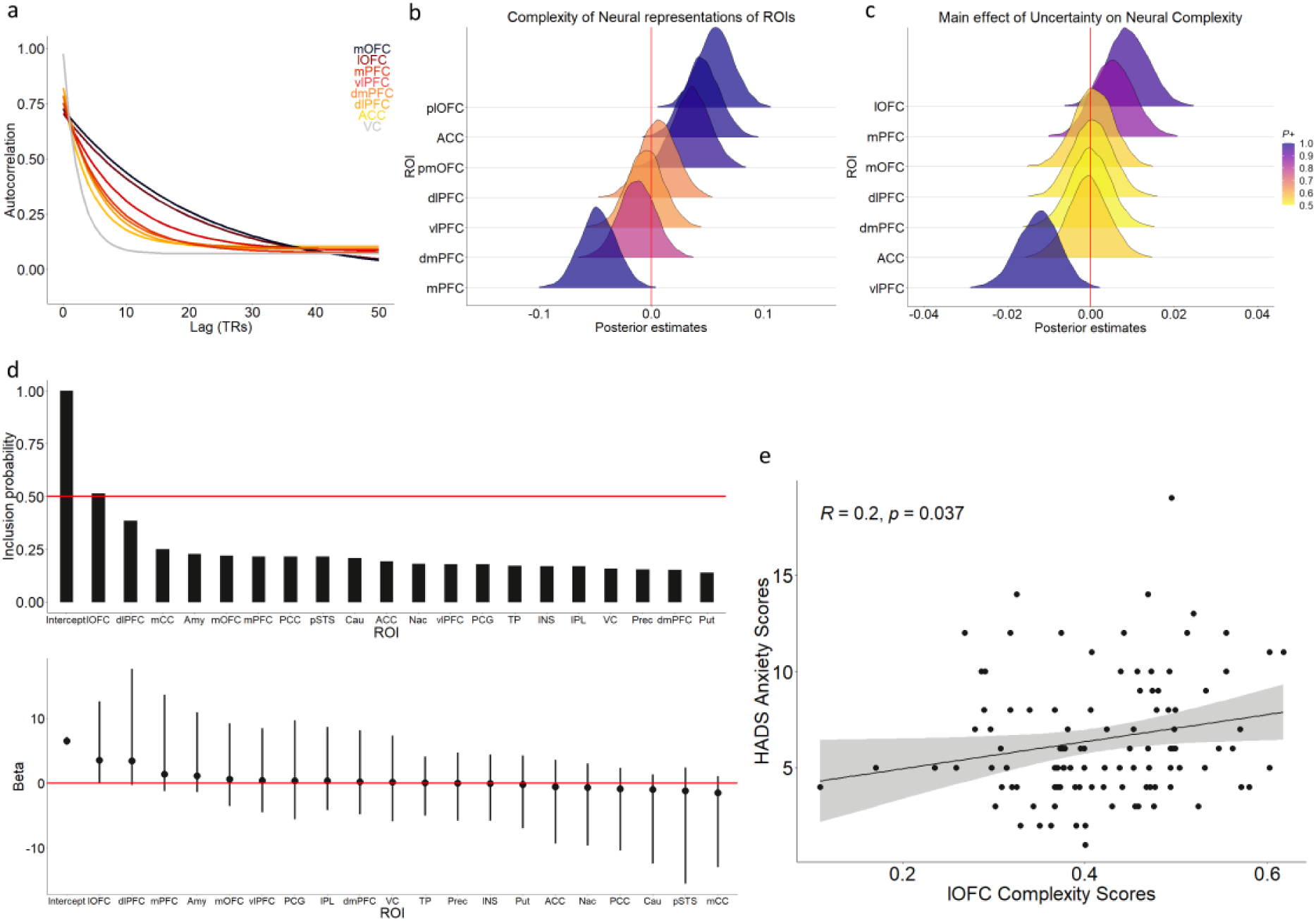
A Hierarchical representation of Uncertainty within the prefrontal cortex. (**a**) Autocorrelation structure of lOFC, mOFC and other prefrontal ROIs along with VC. Each line indicates an exponential function fit to the median of the voxel-wise time series of an ROI across the whole movie (up to 50TR lags). A gradual change (e.g., for mOFC) reflects the rate of integration of information over longer timescales. (**b**) Posterior density estimates of representational complexity among the prefrontal subregions with lOFC having the highest. (**c**) lOFC is the only region with significant evidence (P = 0.96) for its representational complexity increasing with Uncertainty. (**d**) Bayesian multiple regression. lOFC was the only region with a marginal posterior inclusion probability more than 0.50 (top). Beta estimates of all ROIs (bottom). (**e**) Pearson correlation between the (scene averaged) lOFC complexity scores with Hospital Anxiety and Depression scores (HADS) across participants (n = 109). The shaded region indicates the confidence interval.

#### Neural Complexity

Next, we set out to investigate whether this hierarchy is defined by the underlying complexity of representation of uncertainty. The naturalistic nature of the stimulus captures varying levels of uncertainty in the movie, possibly entailing a hierarchy of representational complexity of uncertainty. To estimate this, we first quantified neural complexity adopting a dimensional-reduction approach reported recently ^58^. We performed PCA on voxel-wise data of each Ascent for each subject. We hypothesised that with an increase in representational complexity, the number of PCs required to explain 90% of the variance of signal would increase. Thereafter, we deployed a hierarchical Bayesian model to assess whether the neural complexity of ROIs scaled with uncertainty. We used the same model architecture as our earlier analysis, taking only Ascent timepoints. This enabled us to obtain a robust estimate of regions showing significant evidence for representational complexity and the effect of uncertainty modulating this complexity (Fig. 6b<u>, c</u>). The results revealed that lOFC (lOFC: β_mean_ = 0.06, 95% HDI = [0.02, 0.09], P = 0.99) and mOFC (OFC: β_mean_ = 0.04, 95% HDI = [0.003, 0.07], P = 0.97) along with ACC (ACC: β_mean_ = 0.04, 95% HDI = [0.01, 0.08], P = 0.99) showed substantial evidence for higher complexity values (Fig. 6b). Crucially, we found that lOFC was the only ROI which showed evidence for being positively modulated by uncertainty (β_mean_ = 0.009, 95% HDI = [-0.0006, 0.02], P = 0.96) (Fig. 6c).

In summary, our analyses show that the prefrontal areas exhibit a hierarchical ordering in processing temporally evolving uncertainty in affective experiences with the mOFC and lOFC at the highest level. This hierarchy is reflective of the underlying representational complexity of the regions mediated by uncertainty.

Combining the results so far, lOFC appears to be the only area sensitive to uncertainty (evident from the change in the activity), contributing to a lower-dimensional neural signal underlying the temporal uncertainty and having the largest representational complexity that scales in response towards changing uncertainty. These results, combined with the connectivity dynamics of these regions, adds a new dimension and substantial evidence to this region’s role in processing and responding to emotion-related uncertainty in real-world scenarios.

### Complexity scores of lOFC can predict baseline anxiety scores

A key role of lOFC reported from multiple studies is its ability to predict future outcomes ^52,59,60^. One possibility is that the representational complexity is driven by its role in predicting future outcomes in an emotional situation. Given the evidence of lOFC implication in anxiety disorders ^30^ and its role in negative stimulus-outcomes, we speculated that individuals with highly complex representations would be predicting more adverse outcomes, which would reflect in their baseline anxiety levels. The key assumption here is that these individuals would naturally react to similar real-world situations, thereby having higher baseline anxiety scores. To test this, we conducted a Bayesian multiple regression (using Bayesian model averaging) with the scene-averaged complexity values of each ROI being predictors and HADS anxiety scores as the dependent variable. Results showed lOFC to be the only significant predictor as reflected by its marginal posterior inclusion probabilities (pip) being greater than 0.5 (β = 3.52, HDI = [-0.01, 12.6]) (Fig. 6d). Participant anxiety scores significantly correlated with lOFC complexity levels (r = 0.2, p = 0.037) (Fig. 6e). This sheds new light on what could be a prominent role of lOFC in affective disorders – estimation of uncertainty and the misrepresentation ^30^ of which can lead to being at-risk for developing anxiety and related disorders.

### The emergence of uncertainty from valence in a hierarchical Bayesian model

The results so far show evidence for uncertainty as a defining parameter in mediating complex emotional dynamics. It should be noted that we are certainly not disregarding the role of valence in emotions as suggested by decades of study, despite differences in the literature ^8,9^. Our definition of uncertainty– emerging in situations with different values of the possible outcomes - suggests that uncertainty can be computed from valence, more specifically the changing valence in time across the movie, as a continuous inference. This can establish a relationship between these two parameters previously unknown in affective dynamics literature.

The key feature of Bayesian inference is estimation of uncertainty which led us to hypothesize that during affective experience, this uncertainty is possibly evaluated in a continuous Bayesian inference. Moreover, a hierarchical scheme can be an optimal way to implement this in the context of socially complex temporal emotions. To investigate this, we used a hierarchical Bayesian learning model (HGF, see Methods) to simulate belief trajectories of a (Bayes-optimal) agent (Fig. 7a), having access to the valence information of the movie. Our main interests were threefold a) whether subjective uncertainty can naturally emerge from ongoing valence in the movie, b) if this required a model architecture like what we observed in our neural results and c) whether there is any agreement between model-based estimates of (hidden) states of uncertainty and neuronally observed states, and to this end we made two assumptions. First, this Bayesian ideal observer was defined in a way to minimize the surprise about the input. Secondly, we gave the valence ratings as input to the agent, hypothesizing that the temporal changes associated with valence fluctuation (reflecting the dynamic nature of content in the scenes) would lead to an overall estimation of its volatility, which would be reflective of the (participants’) uncertainty. HGF, thus models the belief trajectories of this Bayesian ideal observer, based on the temporal evolution of the movie valence.

**Figure 7:**
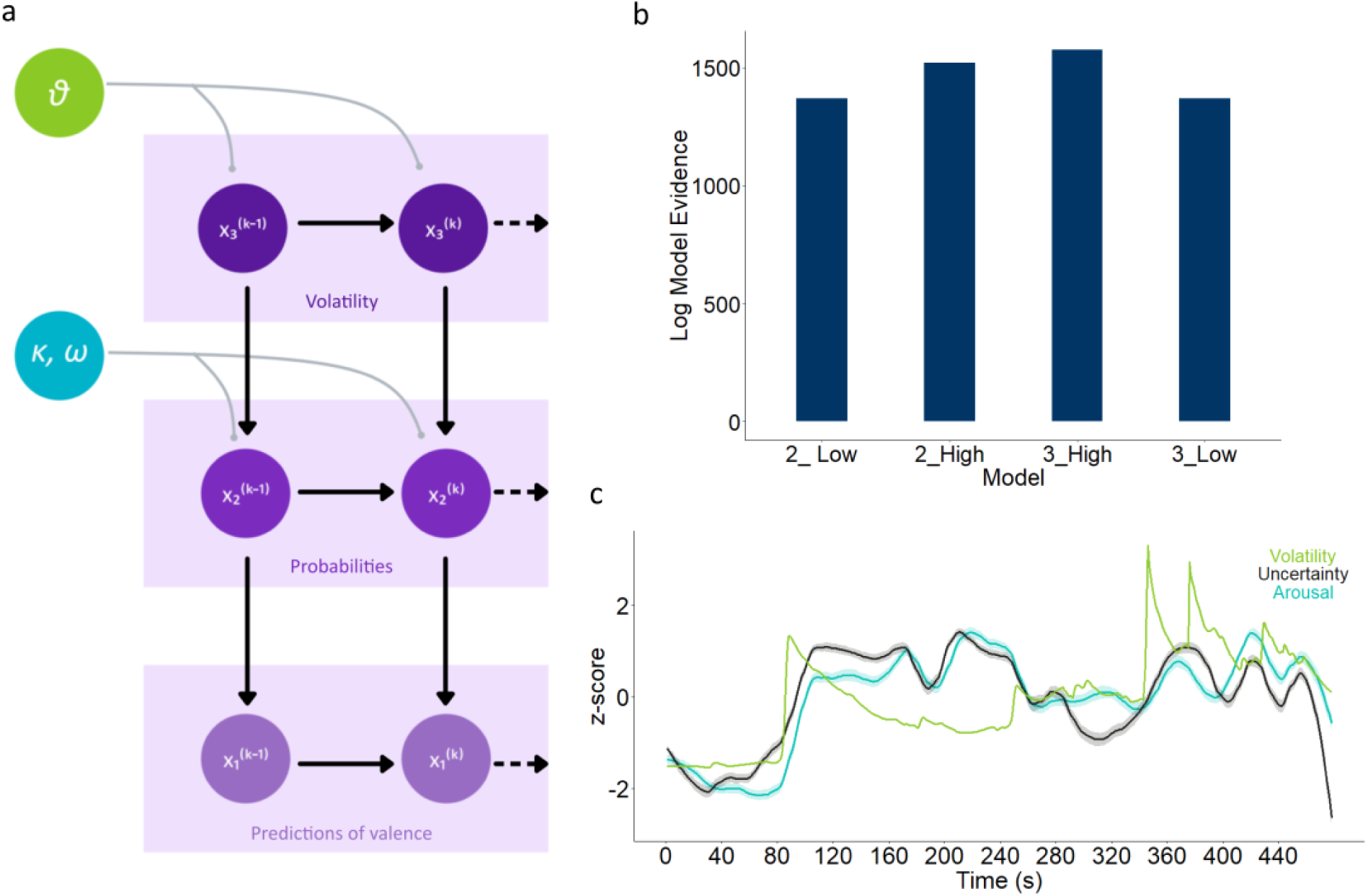
Uncertainty naturally emerges from Volatility of Valence in a generative model. (**a**) Graphical depiction of the Hierarchical Gaussian Filter. Beliefs about the information on Valence is estimated at the level x_1_, which is based on predictions from x_2_, the level above. x_2_ itself varies as a Gaussian random walk, whose step-size is determined by x_3_, which estimates the overall volatility related to the valence time course. The levels themselves are coupled via a phasic, time-varying parameter kappa (***κ***), and a tonic, time-invariant parameter omega (***ω***). x_3,_ in addition, is only governed by theta (***ϑ***). Together, these parameters outfit the step size and the resulting belief trajectories at each level. (**b**) Based on previous neural results, four models were compared - A 2-level and a 3–level HGF with low and high coupling between the levels. 3-level HGF with high coupling (3_High) was the winning model with the highest log model evidence. (**c**) Belief trajectories (z-scored) of Volatility of valence (x_3_) was found to correlate significantly with Uncertainty (r = 0.46) and Arousal time (r = 0.5) courses.

We deployed 4 variants of agents motivated by our neural findings. Specifically, the hierarchical nature of processing uncertainty and high functional connectivity within the prefrontal cortex, with a principal role of the lOFC, led us to test 4 competing (Bayes-optimal) agents by changing the prior parameters of each. The defining properties were number of levels and coupling strength between the levels, reflecting the neural results. This was realized by having 2-level HGF and 3-level HGF models, each with high and low coupling between the levels. Therefore the 4 agents were 2-HGF with low coupling, 2-HGF with high coupling, 3-HGF with low coupling and 3-HGF with high coupling. Model inversion was conducted by variational inference and model adjudication was done by selecting the one with the highest log-model evidence (LME) (Fig. 7b) (Supplementary Table 2). The winning model was 3-HGF with high coupling, mirroring neural results.

We found that, belief trajectories of the Bayesian agent’s third (final) layer, which signifies volatility estimates of the movie valence, was significantly correlated with both subjective ratings of Uncertainty (r = 0.46, p =1.62E-26) and Arousal (r =0.5, p = 4.529E-32) (Fig. 7c). This result and the fact that the 3-HGF High coupling model had more correlation with uncertainty ratings than other competing models (Supplementary Table 2) suggest that a Bayesian hierarchical inference scheme can optimally give rise to estimates of uncertainty from valence. To uncover our last inquiry, which is to test whether there is an agreement between the model-based estimates of uncertainty and that inferred based on neural state estimation, we conducted an HMM for extracting the neural states of uncertainty.

### Neural states of uncertainty in the lateral OFC are consistent, specific, and temporally extended

The temporal nature of ongoing affective uncertainty coding in the lateral OFC suggests that the underlying states might be transitioning from periods of stable periods of uncertainty to highly unstable uncertainty, and back. We used a multivariate HMM, taking individuals as an independent sequence to uncover such states from the lOFC. This generative modelling had a few assumptions. First, even though there might be differences among participants in their experience of the uncertainty, they would visit similar states in the movie. Second, mirroring our previous results, different areas should show these states differently with regions sensitive to uncertainty showing more robust signatures than others. We fit a few models for each ROI with pre-specified states ranging from 2 to 10. Model adjudication was done by selecting the one with the least Bayesian Information Criterion (BIC) (Supplementary Table 3).

The output showed 5 distinct states in the lateral OFC (Fig. 8c). State 4 was specific to the control segment in the movie (state 4, variance of Uncertainty = 13%). State 5 and State 3 were seen in periods of highly fluctuating uncertainty (State 5: variance of Uncertainty = 26%, State 3: variance of Uncertainty = 38%). State 1 and 2 were observed in periods of relatively sustained periods of uncertainty (State 1: variance of Uncertainty = 16%, State 2: variance of Uncertainty = 14%). It is interesting to note that Ascent and Descent segments were found in State 3 and State 5, suggesting how these states might be mediating the ongoing emotional experience.

**Figure 8:**
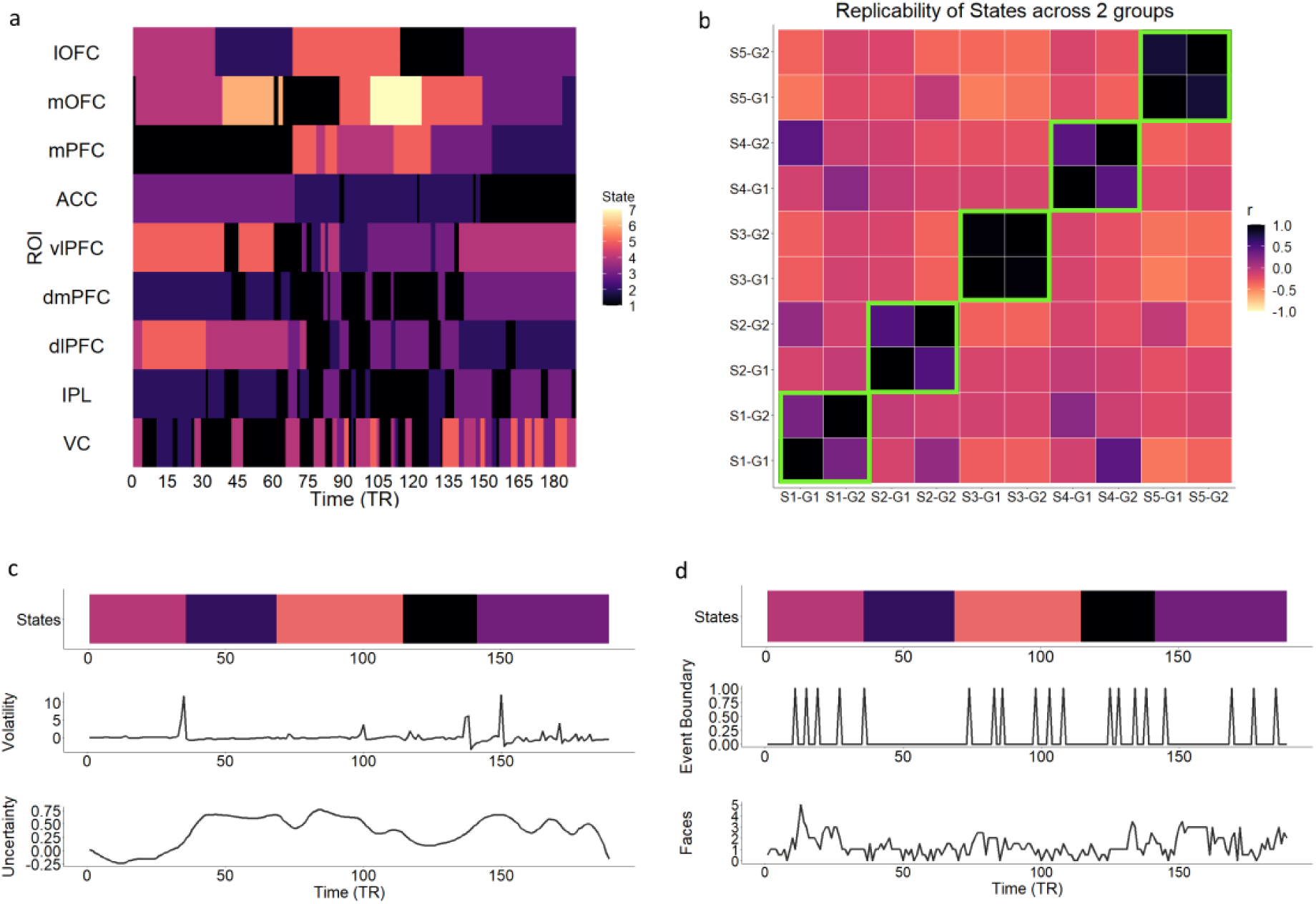
Neural states of Uncertainty within the lOFC are consistent, specific and temporally extended. (**a**) Sequence of neural states with lOFC having sustained and more persistent states. Based on Bayesian information criteria (BIC), different ROIs exhibited different number of optimal states (n) - ACC, IPL and dmPFC: n= 3; VC, vlPFC, dlPFC, mPFC, lOFC: n= 5; mOFC: n = 7. (**b**) lOFC states show high replicability across groups when the sample was randomly split into two subgroups (n_1_=50, n_2_=61). (**c**) lOFC states are specific to ongoing volatility updates (estimated from HGF) as well as to states of fluctuating Uncertainty. (**d**) lOFC states are not due to uncertainty related to faces within the movie nor due to event transition related contextual uncertainty.

HGF models belief trajectories through time, and it promptly updates whenever errors in future prediction of valence estimates occur. These result in updates to ongoing predictions, which is also a function attributed to OFC ^59,61^. In order to test whether the behavioral consistency was observed neurally, we compared the HMM state changes of lOFC with the belief updates of predictions in the Bayesian model. We found out that all the HMM state transitions coincide temporally (on average within <3 time points) of the belief updates estimated from the model (Fig. 8c). Although there are other belief updates unaccounted for by the lOFC, the reverse wasn’t true. lOFC states, in addition, were specific to the dynamic uncertainty, and not associated with detection of faces or event boundaries (Fig. 8d). We did this analysis as the OFC is shown to encode social information off faces ^62^ and (sudden) perceptual event transitions might cause a temporary spike in uncertainty. Regardless, it showed that these states were specific to the uncertainty dynamics, and not to purely perceptual sources of uncertainty.

Given the distributed nature of uncertainty processing and the differential role of different regions to it, we compared the neural states of uncertainty of lOFC with other regions. The key pattern that emerged was a temporally persistent nature of the states within lOFC compared to other regions. The PFC in general elicited this pattern of temporally extended states more than other areas involved in perceptual processing like IPL and VC (Fig. 8a).

A replication analysis was also conducted by splitting the sample into two groups with 50 and 61 participants, and aligning the resulting states by the Hungarian algorithm (which maximized spatial similarity between the states by the Pearson correlation) showed all 5 states were significantly correlated between the groups (Fig. 8b) (State 1: r = 0.30 , p = 2.41E-05 ; State 2: r =0.50 , p = 2.81E-13 ; State 3: r = 0.97, p = 1.08E-111 ; State 4: r = 0.45, p = 6.80E-11 ; State 5: r = 0.8 , p = 8.74E-44). Moreover, these states were also consistent across individuals when we conducted an individual level HMM for lOFC and aligned it with the Hungarian algorithm (Supplementary Fig. 6).

Our findings conclude that firstly, the lOFC tracks the subjective uncertainty by updating ongoing beliefs about this uncertainty evident from its neural states. Second, these states are specific to uncertainty due to high level belief changes rather than low level sources of uncertainty, essentially showing a predictive code for naturalistic complex emotions within the prefrontal cortex.

## Discussion

In our present study, we put forward the notion that uncertainty is a defining dimension of emotion dynamics. By incorporating Bayesian modelling on a complex naturalistic stimulus, we found that uncertainty has a significantly larger modulatory effect and explanatory power on affective experience compared to valence. This unites emotion dynamics with theoretical grounds of predictive coding ^63^ and Bayesian learning ^28,64,65^, where uncertainty acts as the inferential parameter. Moreover, this provides supporting evidence to works which have shown that valence-based characterization of emotion reliant on a positive-negative axis, is insufficient in explaining complex naturalistic emotions^8^, which are contextual ^66^ resembling real-world affective dynamics. Although the relationship between uncertainty and emotion has been highlighted previously in the literature ^1^, uncertainty as a dimension to study complex emotion dynamics has never been explored. We demonstrated that for both increasing and decreasing affect conditions, uncertainty elicited significantly larger effect in mediating arousal relative to valence.

Converging evidence from both supervised and unsupervised approach suggested that during emotional experiences this uncertainty is mediated by a distributed set of areas in the prefrontal cortex with a central role of lOFC. Apart from the PFC, contributions from parietal, insular and cingulate regions were also evident in the widespread activation. It is interesting to note that a recent study^41^ showed similar widespread neuronal networks were responsible in processing fear rather than dedicated fear centres as was previously thought. Our study suggests that this might be true for other emotional experiences, not specific to fear processing and could be a fundamental mechanism for emotions in general. The increasing PFC activation in response to uncertainty as observed from the model was also evident when carried out in an independent, unsupervised analysis, aligning with literature showing this regions’ heavy involvement in tasks with uncertainty ^26,36,39,42^. This was apparent in a low dimensional neural signal which varied strongly with uncertainty, reflecting the Ascent phase of Arousal. Since affective experiences are temporally dependent, this unsupervised approach enabled us to investigate the temporal evolution of the (lower-dimensional) neural signals of our stimulus to uncover distinct spatiotemporal neural signals tracking the evolving uncertainty, adding credence to our hypothesis. Crucially, a consistent observation that emerged from both the analyses was the opposing behaviours exhibited by the lateral and medial OFC, suggesting a functional dissociation.

The role of OFC has been implicated extensively in predicting outcome value ^52,59^ as well as representing task space ^67,68^. The importance lies in its central role in estimating the future by representing the (present) latent state, which is fundamentally dependent on accurate representation of uncertainty of the current and future states. In a dynamic setting, this demands rapid representation of changing uncertainty to estimate future states. Additionally, OFC possesses dense bidirectional connections with midbrain dopamine neurons, the activity of which is scaled by outcome uncertainty^13,69,70^. This strongly suggests a key role of OFC in mediating temporally complex real-world emotional experiences, which has not been studied previously. Thus, we focused on investigating distinct functional network dynamics within the OFC in temporal processing of uncertainty, which has not been explored previously. We found strong coupling of lOFC with lateral fronto-parietal regions (but not DMN) likely driving the top-down predictions to direct attentional processes in response to exogenous uncertainty. A contrasting pattern was observed for mOFC which coupled strongly with DMN possibly reflecting the subjective ^57^ processing of uncertainty with relation to present states, past experiences or future simulations. Thus, our findings demonstrate a functional double-dissociation within OFC highlighting the differential but ultimately coordinated role of lOFC and mOFC in complex emotional experiences. Future studies can be set up to help identify more nuanced functional differences in this division of labour between the medial and lateral sectors of OFC in emotion ^27,71,72^.

Different connectivity strengths between different regions of PFC suggests that the difference might exist in encoding varying levels of complexity of uncertainty. We indeed found a hierarchical processing of uncertainty within the prefrontal cortex with mOFC and lOFC occupying the highest level. These regions showed varying timescales of integrating information suggesting that encode varying levels of complexity. A growing body of evidence ^73,74^ demonstrates hierarchical processing in brain networks, and this adds a novel addition to it by showing such an organization for representing uncertainty, temporally as well as spatially with OFC at the highest level. By quantifying the representational complexity adopted from a recent work ^58^, we found that lOFC appeared to encode the highest level of complexity mediated by uncertainty during increased arousal. An advantage of this approach is purely data-driven in nature than being the output of a prespecified model. A key response to outcome uncertainty is representing the future states, and consequently the expected value or free energy ^75^. Hence, a deeply uncertain situation demands a more complex representation of the outcomes. The specificity of lOFC to respond to uncertainty by increasing its complexity and activation outlines a dedicated role evolved for this.

On a computational level, we found that the volatility belief trajectories obtained from Bayesian generative modelling correlated with the uncertainty. This posits that uncertainty can naturally emerge from the change in valence in a hierarchical and highly coupled architecture. Moreover, the trajectory estimates of highest level aligning with neural states in the lOFC points to an account of a predictive code in the service of emotional experiences. The long temporal duration of information integration in the lOFC is further substantiated in its latent state which was persistent as well as distinct towards the nature of uncertainty. Specifically, unlike other regions like VC, IPL, the latent states in lOFC remained relatively stable in the face of fluctuating uncertainty. These results suggest that during real world emotional instances, these higher-order regions encode affective state estimates and monitor the variance of these estimates concurrently. It is important to acknowledge that it remains hard to differentiate whether the representing uncertainty is a response to an environment or an intrinsic subjective dimension (especially in hierarchical systems where uncertainty in one level can influence the next). However, the neural complexity responses in lOFC being predictive of a subjects’ baseline anxiety scores is evidence of how it could be a combination of both, with overrepresentations affecting emotional well-being. In summary, lOFC representations were highly complex, modulated by uncertainty, is predictive in nature and showed a functionally distinct processing compared to the mOFC. These results broadly underwrite the fundamental nature of neural uncertainty representation in emotions and advance our understanding on the massive prevalence of OFC in affective disorders.

Although, this movie has been used in a wide range of studies evincing its rich content, further research is required to ensure the generalizability in explaining different degrees of complex emotions. The dominating negative valence and absence of truly positive valence in our stimuli might invite criticism on establishing uncertainty as an explanatory construct for emotions. However, this portrays a necessity of a wider spectrum for valence (i.e., highly positive) as compared to uncertainty. One can safely imagine nonetheless that uncertainty can influence positive emotions strongly as well ^76^ (e.g. spoilers reduce the intensity of experiences even for positive outcomes).

Explicitly modelling the dynamic affective experience using a movie enabled us to richly sample real-world emotions and quantify the uncertainty in estimating emotional outcomes. We formalize a specific role for the lateral OFC in representing and responding to uncertainty in socioemotional settings, in a predictive manner, with a general role for the prefrontal cortex which processes this in a hierarchical organization. We hope that incorporating uncertainty might help uncover many aspects of complex emotions which are difficult to reconcile with a purely two-dimensional or discrete approach. Bayesian brain frameworks have contributed immensely to our comprehension of various cognitive functions, but in future its most impactful effect would be on understanding the most unpredictable of them all – our emotions.

## Methods

### Stimulus

We examined the behavioural and neuroimaging responses of participants to a tailored version of a black and white movie by Alfred Hitchcock called “Bang! You’re Dead!” The entire 25 minutes movie was compressed into an 8-minute movie clip maintaining the original sequence (context) of the movie ^77^. This movie has been widely used as a stimulus in various cognitive domains validating its strength as a robust naturalistic stimulus.

### Behavioural participants

We used an online data collection platform Prolific (https://prolific.co/), to collect the behavioural data from 3 independent groups of participants, demographically matched with the fMRI participants (UK). They were further screened based on their educational qualification and mental health status to reduce potential confounds. 60 participants (20 participants per group) were recruited for this study with ages ranging from 18-30 yrs. All participants were right-handed, native English speakers and had normal or corrected-to-normal vision and hearing. The participants reported no previous history of watching this movie.

### Behavioural design

The stimulus was presented to the participants on custom software with a vertical slider on the side (outside the movie frame), with encoded values from -1 to 1. Each participant was instructed to use headphones, pay attention, and continuously rate the movie while watching it, using their mouse to move the slider up and down. Ratings for valence, arousal and uncertainty were acquired from 3 separate groups. The study was approved by the Institutional Human Ethics Committee of the National Brain Research Centre, India (IHEC, NBRC).

### Behavioural analysis

We computed behavioural intersubject correlation (bISC) to identify reliable behavioural responses using the leave-one-out approach across 20 subjects for each group and removed subjects with less than r = 0.2. This yielded a total of 50 participants (Arousal – 17, Uncertainty -16, Valence – 17) (Mean age = 24.5, SD = 3.75, 27 female).

The behavioural time courses were z-scored for each subject and averaged to obtain a mean time course for each behavioural measure. Each averaged time course was smoothed using local regression smoothing (*loess*) with a span of 21 TRs. *Loess* is a non-parametric approach, particularly suited to variable time courses bounded within a range (here -1 to 1). The same span was used to calculate sliding-window intersubject analyses later ^78^. To isolate the emotionally dynamic segments, we identified the local maximas and minimas from the mean normalised and smoothed Arousal rating, using the *findpeaks* and *islocalmin* function in MATLAB, respectively. We specified the emotionally intense anticipatory phase as Ascent (trough to peak segments) and the release phase as Descent (peak to trough). We only took the corresponding Ascents and Descents with a minimum of 6 timepoints for further analyses.

We excluded the peak timepoints depicting the outcome from these segments. The Control segment was specified as the timepoints where Arousal, Valence and Uncertainty followed a flat trajectory with less than 1 SD from the baseline. A total of 1 Control, 5 Ascent and 5 Descent segments were identified for subsequent analyses. Additionally, the behavioural time courses were downsampled to match the BOLD time course.

### fMRI participants

A total of 135 participants between the age of 18-35 years were selected from the healthy population-derived Cam-CAN cohort ^79^. Following data preprocessing, a sub-sample of 111 participants (mean age = 28.5, SD = 4.93, 63 female) were selected for the fMRI data analysis. All participants of this cohort were assessed for general cognitive functions across multiple domains like attention, executive control, language, memory, emotion, action control and learning and had an MMSE score>24. All participants were native English speakers. We did not exclude participants based on handedness, given the large sample size. Written informed consent was obtained in accordance with the Cambridgeshire Research Ethics Committee.

### fMRI acquisition

The fMRI scans were carried out using a 3T Siemens TIM Trio with a 32-channel head coil. High-resolution structural images were acquired using T1-weighted magnetisation Prepared Rapid Gradient Echo (MPRAGE) pulse sequence (Repetition Time (TR) = 2250 ms; Echo Time (TE) = 2.99 ms; Inversion Time (TI) = 900 ms; Flip Angle = 9°; Field Of View (FOV) = 256 × 240 × 192 mm; voxel size = 1 mm isotropic). The functional scans comprised of resting-state and movie watching sessions. For this study, we used the functional data from movie-watching session, which were collected using a T2*−weighted Echo Planar Imaging (EPI) pulse sequence (TR = 2470 ms; TE = 9.4 ms, 21.2 ms, 33 ms, 45 ms, 57 ms; Flip Angle = 78°; 32 axial slices (acquired in descending order); slice thickness = 3.7 mm; FOV = 192 × 192 mm; voxel size = 3 × 3 × 4.44 mm) ^77,79^. The acquisition time (TA) for this session was 8 min 13 s, which resulted in 193 functional volumes.

### fMRI preprocessing

Neuroimaging data were preprocessed in MATLAB (2019b) using Statistical Parametric Mapping (SPM12) (https://www.fil.ion.ucl.ac.uk/spm). The preprocessing was performed on raw signals and included spatial realignment, slice-timing correction, coregistration of functional (T2) to anatomical (T1) scans, affine transformation to MNI template (MNI152), resampling to 2mm isotropic voxels, and spatial smoothing with a 6mm x 6mm x 9mm full-width at half-maximum (FWHM) Gaussian kernel. Six motion parameters (inter-scan X-, Y-, Z-displacement and pitch, roll, yaw parameters) were regressed out from functional data by least square regression. Subjects who had maximum framewise displacement greater than 1 mm or 1.5° were discarded from further analysis. Following preprocessing, the first 4 functional scans were discarded, which yielded a total of 189 scans for each subject. Thereafter, all functional data underwent voxel-wise detrending and was band-pass filtered in the range of 0.01-0.1 Hz using a second-order Butterworth filter.

### Whole-brain Spatiotemporal ISC

To calculate the shared neural engagement ^80^ of the CamCan participants, we performed a whole-brain spatial ISC on the neural data using Brain Imaging Analysis Kit (http://brainiak.org). First, we extracted the whole-brain BOLD signal for each subject using the Power Atlas ^45^ which consists of 264 cortical and subcortical parcellations. Thereafter, we computed a dynamic leave-one-out ISC using a sliding window ^44^ of 21TR with 1TR overlap by correlating the BOLD signal of each subject with the mean BOLD signal of the rest for each window and for each region of interest (ROI). The Pearson correlation coefficients were averaged for each window, Fisher-z-transformed and averaged across subjects ^81^ which resulted in a single time-course of mean ISC across the whole movie. Since we did not have any behavioural measure of their subjective experience, the dynamic ISC enabled us to capture their shared response ^81^ varied in time. This was further correlated with the downsampled mean Arousal time-course of the behavioural group to assess the similarity between behavioural and neural responses.

To investigate the shared response across three conditions, we also performed a temporal leave-one-out ISC for Ascent, Descent and Control by extracting the specific time points for each Ascent and Descent and concatenating them across time. This yielded three matrices (m x n x p), one for each condition, with m = 264 ROIs, n = TRs and p = 111 participants. The resulting p correlations were averaged to get the mean ISC for each condition. The same method was extended to calculate scene-wise ISC for Ascent and Descent. Bayes factor (BF) analysis was done on the paired *t-tests* between Ascent and Descent with Control ^82^ (using a default Jeffreys Zellner Siow (JZS) prior) using the *BayesFactor* package in R.

### Neural General Linear Model (GLM) Analyses

To identify the neural activity corresponding to the emotionally dynamic conditions, we modelled the entire movie viewing BOLD time-course by fitting a GLM which included discrete regressors for each Ascent, Descent and Control segment identified from the independent Arousal ratings. Six head-motion parameters defined by the realignment were added to the model as nuisance regressors. Each regressor was convolved with a canonical hemodynamic response function, and the resultant GLM produced 11 regressors of interest (5 Ascent, 5 Descent and 1 control).

Since our hypothesis was specifically on the Ascent condition, linear contrast images for each Ascent were created by subtracting the parameter estimate of Control from each Ascent (Ascent>Control). A group-level random-effects analysis was conducted using the individual contrasts as input. We performed a whole-brain one-sample *t-test* to identify voxel-wise activations for each Ascent. Thereafter, we thresholded the statistical maps at q < 0.05 FDR with an extent threshold of 25 voxels (k = 25) which was performed using NeuroElf (http://neuroelf.net/).

### ROI definition

Due to the dynamic nature of the naturalistic stimulus in this study, each Ascent cannot be treated as a trial of the same condition. Hence, we defined our ROIs from activation clusters present in at least 3 (out of 5) Ascents. The logic for this selection process was to include the regions that might not survive a conjunction analysis of the Ascents but showed significant activation (k > 100) in the majority of the Ascents. After analysing the contrast maps, we selected the following ROIs for our initial analyses – dorsolateral prefrontal cortex (dlPFC), dorsomedial prefrontal cortex (dmpFC), ventrolateral prefrontal cortex (vlPFC), medial prefrontal cortex (mPFC), lateral orbitofrontal cortex (lOFC), medial orbitofrontal cortex (mOFC), anterior cingulate cortex (ACC), Amygdala (Amy), posterior cingulate cortex (PCC), precuneus (Prec), precentral gyrus (PreCG), visual cortex (VC), caudate (Cau), anterior Insula (INS), inferior parietal lobe (IPL), nucleus accumbens (Nac), putamen (Put), temporal pole (TP), posterior superior temporal sulcus (pSTS). We defined ROI masks from Brainnetome Atlas or constructed spherical ROI masks using peak coordinates after analysing the extent of activation clusters. We extracted the BOLD and the percentage change signal of these ROIs using MarsBar.

### Bayesian Hierarchical Regression

To delineate the effects of Uncertainty and Valence on emotion dynamics, we employed a Bayesian Hierarchical Regression model ^49,83^ rather than the frequentist approach using conventional “p-value” statistic with binary significance, which has been shown to constrain inferences ^84^. This enabled us to have a spectrum of inference with necessary statistical rigour ^85,86^ taking into account the complexity of the data. The way we constructed the model (Fig. 2a) incorporates the effects of Uncertainty and Valence as their interaction with Arousal (denoted categorically as Ascent/Descent), simultaneously including the variability within-subjects and across subjects on the level of ROIs in one single multidimensional model. Uncertainty and Valence were parametrically fed into the model as the averaged values of each segment of Ascent and Descent, thus addressing scene-specific biases. Therefore, this model incorporated individual subject variability (subject level variance), content variability (having scene-specific values for Uncertainty and Valence), ROI level variability (having a varying slope and intercept over each ROI) and participant dependent variability on Uncertainty/Valence/Arousal (varying respective slopes for each subject).

The dependent measure of the model input was given by the % BOLD signal for each of the 20 ROIs for each Ascent or Descent relative to Control and was modelled as a Student t-distribution -

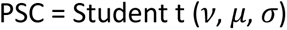

where *v* is the degrees of freedom, *σ* is the scale parameter of the distribution and *μ* is the mean which can be expressed as

*μ* = α + α_j_ + α_i_ + β_1_. Arousal:Uncertainty + β_2_.Arousal:Valence + γ_1j_.Arousal:Uncertainty|ROI + γ_2j_.Arousal:Valence|ROI + γ_3i_.Arousal|Participant + γ_4i_.Uncertainty|Participant + γ_5i_.Valence|Participant + *σ*_i_ + ε

Here, α is the intercept, α_j_ is the varying intercept over *j* ROIs, α_i_ is the varying intercept over i participants. β and γ reflect the slope and varying slope of the predictors, respectively. β_1_ represents the slope for the cross-level interaction term of Arousal and Uncertainty. β_2_ denotes the slope for the cross-level interaction term of Arousal and Valence. It should be noted that Uncertainty and Valence are continuous predictors (which are mean-centred), while Arousal is a two-level categorical predictor (Ascent/Descent), which when unpacked would lead to Ascent:Uncertainty, Ascent:Valence, Descent:Uncertainty and Descent:Valence where ‘:’ denotes the cross-level interaction. γ_1j_ indicates the varying slope term where the cross-level interaction term (between Arousal and Uncertainty) varies across all the ROIs. Likewise, γ_2j_ denotes the varying slope of Arousal and Valence interaction over the ROIs. γ_3i_, γ_4i,_ and γ_5i_ are varying slopes where each of the three predictors are allowed to vary across all i (=111) participants. σ_i_ and ε connote subject level and population-level variance terms, respectively. Weak, noninformative priors were applied over the model terms.

- *α* ∼ *Student_t* (3, 0.5, 2.5)
- *β* ∼ *Student_t* (3, 0.5, 2.5)
- *γ* ∼ *Student_t* (3, 0.5, 2.5)
- *σ_j_* ∼ *Half Student_t* (3, 0, 2.5)
- *ε* ∼ *Half Student_t* (3, 0, 2.5)

The model output gives a full posterior distribution than point estimates. Posterior estimation was done using the *brms* package in R ^87^, through Hamiltonian Monte Carlo for 2 chains of 4500 samples with the first 1000 discarded from each. Model convergence was assessed through R-hat statistics which were found to be 1.00, sufficiently large, estimated sample size (ESS) for stable posterior estimates and visually inspecting the chains for convergence and large autocorrelations. Posterior predictive checks for model validation were conducted by simulating 1000 samples from the posterior and fitting with the empirical data (Supplementary Fig. 3). Analysis was focused on two levels – the first level estimated the main effect (population-level effects) of Arousal: Uncertainty and Arousal: Valence interaction term on the neural activation. This helped us distinguish between the main effects of Uncertainty and Valence. The second was on group-level, which was performed on the level of ROIs capturing the effect of Uncertainty and Valence on ROIs separately. Significance levels, or evidence in this case, were performed by estimating the evidence of posterior probability. Evidence of P > 95% is equivalent to the frequentist depiction of significant p-values. In addition, we also quantified the effect size by computing the Bayes factor (calculated by the Savage Dickey ratio).

We also ran a null model (without Uncertainty and Valence), a Valence only model (ValM) and an Uncertainty only model (UncM). This was done to understand the effect of each of these two parameters in explaining the model. The null model assumed the neural activations to be explained through only the phase of emotions (i.e., Ascent or Descent), and the Combined model considered influence of both Uncertainty and Valence. These models were assessed by their R^2^ and expected log predictive density (ELPD) values (relative to the null model) to explain how each of them fits the data.

### Spatiotemporal PCA

To capture the lower dimensional neural signals corresponding to Ascent and Descent we performed a spatial principal component analysis (PCA). We averaged the extracted BOLD time courses of 20 ROIs for each condition (Ascent and Descent) and concatenated the mean Ascent (10 TR) and mean Descent (10 TR) BOLD data for each subject which yielded a p x n matrix (p = 20 TR, n = 20 ROI) for each subject. After column-wise centring and scaling by (n-1), we performed the PCA computed with a covariance matrix of the p x n matrix, using singular value decomposition ^88^:

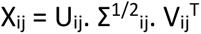

where U and V are eigenvector matrices for the column space (XX^T^) and row space (X^T^X), respectively, while E represents the diagonal matrix with hierarchically arranged eigenvalues. In our study, the principal components (PCs) (U_ij_) represent the covariance structure across time and the eigenvalues represent the amount of variance explained by corresponding eigenvectors. We obtained 20 PCs for each subject of which we only isolated the first two PCs for further analyses. PCA was performed using the *factoextra* package implemented in R ^89^. The first 3 components explained ∼80% variance (PC1 – 45%, PC2 – 26%, PC3 – 9%) (Fig. 4e).

To investigate the temporal evolution of these lower-dimensional components, we implemented a temporal PCA ^50,51^ by estimating the time-series of PC1 and PC2. This was performed by calculating the weighted average of the BOLD time-series associated with each respective PCs. The group-level temporal PC (tPC) time course was estimated by calculating the mean of each tPC time series across the subjects. Thereafter, to estimate the relationship between tPC time series and our conditions, we correlated the tPC1 and tPC2 with Ascent and Descent BOLD time series. We also performed a Pearson correlation between the tPC1 and that of the lOFC and mOFC BOLD signal to examine their relationship with low dimensional neural signals.

### Dynamic and seed based ISFC of OFC

We carried out sliding window intersubject functional analysis (ISFC) ^44,78^ for each participant from their BOLD data. After normalising the BOLD signal across subjects, ISFC was calculated between two ROIs by correlating the time series of an ROI from one subject with the mean time series of another ROI of the rest of the subjects. This was performed over a sliding window of 21 TR, with a step size of 1 TR across the movie. Thereafter, we Fisher transformed the Pearson correlation coefficients and obtained the mean ISFC time series between any two ROIs by averaging across subjects.

This ISFC time series was then correlated with the Uncertainty and Valence ratings. Crucially, the ratings were smoothed using the same span of ISFC (i.e., 21TR) and downsampled to match the duration. The correlation value between an ROI pair and a rating time course reflected the strength of the connection between these ROIs and how it varied dynamically as a function of the behavioural measure, effectively outlining the informational integration between these ROIs for processing that measure. ROIs were grouped into three networks Default mode network (DMN) - mPFC, Prec and PCC, Frontoparietal network (FPN) - dlPFC, vlPFC and IPL, and Limbic - Amy, Cau, Put and Nac.

Next, we computed the correlation between each ISFC time series of a network with the Uncertainty and the Valence time series, obtaining 6 *m* x *n* matrices (3 per condition) where each value represented the correlation coefficient between ISFC of an ROI pair with that of the Uncertainty/Valence time series. Multidimensional scaling was applied on each of the matrices and visualised simultaneously using an R package ^90^. In the resulting figure (Fig. 5a, c), each node represents an ROI. The line between the nodes denotes the correlation strength of the ROI-ROI ISFC with the behavioural time course. The clustering of nodes by the MDS reflects the local distance between the ROIs preserved in a lower dimension space.

ISFC of lOFC and mOFC with the networks were estimated for each condition separately by concatenating the time points of Ascent and Descent (i.e., without the window) and averaged the Pearson correlation coefficients obtained from each of the individuals for each network. After that we performed a Bayes Factor analysis on paired t-test between lOFC and mOFC ISFC with each network in both conditions.

### Temporal autocorrelation

We performed the autocorrelation on each voxel’s time series by computing the average correlation of a voxel’s time series with itself shifted by a lag of t steps where t ∊ {1, 2,…,50}^74^. For each ROI, we took the median across all its voxel autocorrelation time series as the representative shape for that ROI. For the final inference on an ROI, the median across participants was taken and thereby fitted with an exponential function using a robust estimator. This was carried out by the Levenberg-Marquardt algorithm (using the *minpack* implementation in R), which minimises the sum of squared errors as

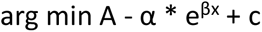

where A is the median autocorrelation for an ROI.

### Representational complexity analysis

We quantified the neural complexity in a two-step procedure. The first step was based on a previous work that used the approach for neural complexity analysis in a concept learning task ^58^. We modified it for our purpose as follows. First, we extracted the voxel-wise activation patterns for each ROI for each of the Ascent. We only limited our analysis to Ascent as we were interested in the effect of uncertainty/valence during the elevated phase of emotional intensity due to rising anticipation. For each participant, we conducted a PCA on the voxel-level spatial patterns for each ROI on each Ascent. The maximum components possible for this (regardless of ROI) was *t* components, the length of the individual scene. The maximum possible components needed to explain 100% of the data was *t* components which suggest maximal complexity. We took the number of components required to explain 90% of the variance as *k*. For example: If ROI_A_ needed only two components to explain 90% variance, while ROI_B_ required 5, one can reasonably imagine the spatial representations of the latter to be much more complex, given a scene. This can be formalised as

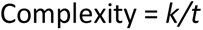

Where *k* is the number of PCs necessary to explain 90% of the variance. Each of the scenes we took had outcome uncertainty; however, specific differences in content can lead to different complexity of representation. For example, a scene where a kid aims a gun at the maid vs aiming it at a father in front of his little daughter would incite different complexities (due to the outcomes themselves), despite the similar underlying uncertainty. To preserve this diversity within the movie, we separately took all of the scenes rather than averaging them.

Secondly, these inputs from different ROIs and across participants were then fed into another Bayesian hierarchical regression model. The model architecture was the same as the one deployed previously except there was no Descent category for Arousal. This ensured robust estimation while incorporating all sources of variability in the stimulus.

Our analyses were focused on the ROI level inference on two aspects - one, which ROIs had significantly more complexity and two, whether the complexity was modulated by the behavioural predictors (Uncertainty/Valence). Although we implemented the model for all the ROIs, we focused the primary analysis on the prefrontal cortex.

### Bayesian multiple regression and HADS

We implemented a Bayesian multiple regression in R using Bayesian Adaptive Sampling ^91^ to predict the hospital anxiety and depression scores (HADS) of the fMRI participants with the ROI complexity values as predictors. HADS were provided by CamCan for all participants. The score ranged from 0 - 21, with scores >8 denoting considerable symptoms of anxiety or depression. We removed 3 participants with 0 HADS scores and thereafter took them as the dependent measure. The averaged complexity values of the ROIs (n=20) were given as the predictors.

Rather than fitting a single regression model, this approach computes multiple models and uses Bayesian model averaging, which gives robust accounting of model uncertainty. Bayes factors are calculated to compare different models and quantify evidence of one model over the other. We used weak noninformative priors: Jeffreys prior was applied over the variance term, *σ* and a (multivariate) Zellner-Siow Cauchy prior was applied over the model coefficients, *β*. A uniform prior was used as the model prior (denoting equal probabilities to all models). Posterior estimation was done by 10000 MCMC samples with the first 2000 discarded.

### Hidden Markov Model

An HMM was fitted to each ROI by taking the individual subject’s voxel-time series as an independent sequence. This creates a single emission sequence for each state across all participants. For each ROI, we fit a number of HMMs from pre-specified states of range (states = [2, 10]). Parameter estimation for the models was done by the Baum-Welch algorithm (a variant of the Expectation-Maximisation algorithm) by iteratively maximising the expected joint log-likelihood of the parameters given the observations and states. These models were then adjudicated by selecting the one with the least Bayesian Information Criteria (BIC) (Supplementary Table 3). Different ROIs had different number of optimal states which had the minimum BIC for the HMM (ACC, IPL and dmPFC optimal number of states = 3, VC, vlPFC, dlPFC, mPFC and lOFC optimal number of states = 5, mOFC optimal number of states = 7). Decoding was done by the Viterbi algorithm by computing the maximum a posteriori state sequence for each of the HMMs.

For the replication analysis for lOFC, we randomly split the total 111 sample size into two subgroups of 50 and 61. HMM was fit to each of the groups by the same procedure. To align the states between each group, we used the Hungarian algorithm ^57,92^, which maximises the spatial similarity between the 5 states for each group. For the single subject replication of states, an individual HMM was fit to each subject, and then state alignment between individuals was done via Hungarian algorithm in the same way as in the group-level HMM.

The number of faces in the movie was detected by feeding the framewise screenshots of the entire movie to a Single Shot Detector (SSD) which is based on a Convolutional Neural Network pretrained on the Caffe Deep Learning kit ^93^. Event boundaries were obtained for the movie from a previous work where 16 participants labelled the event boundaries ^94,95^.

### Hierarchical Gaussian Filter

The Hierarchical Gaussian Filter (HGF) ^96,97^ is a Bayesian learning model wherein an agent learns the temporal evolution of environmental variables as Gaussian random walks. The agent learns this hierarchically, in increasing complexity (as the layers progress upward) based on the uncertainty with which it is able to predict the variables. In other words, this is a generative model that infers how observed inputs are generated by environmental hidden states. Thus, HGF can be construed as predictive coding in the temporal domain, dynamically updating ongoing predictions from prediction errors, and has been extensively used in a range of paradigms like MMN ^98^, gambling tasks ^99^, inferring moral beliefs ^100^ as well as social feedback learning ^101^. The original HGF ^97^ has a perceptual model and a response model, with inputs either binary or continuous.

In our paradigm, since there were no behavioural responses from the scanned participants, we only used a perceptual model on a Bayes-optimal agent using the TAPAS toolbox in MATLAB (http://www.translationalneuromodeling.org/tapas/). On the first layer, x_1_, the evolution of Valence is modelled. The level up, x_2_, estimates the agents’ belief about how the underlying distribution that gives rise to valence changes. Finally, the level x_3_, estimates the overall volatility associated with the movie valence. Whenever a prediction error (i.e., misestimation of current valence from predicted valence) occurs, the layers get updated. This update itself is equivalent to the prediction errors weighted by the precision, ***π*** (inverse of uncertainty). Thus, at each level the prediction errors are weighted by their relative precisions (ratio of how much new learning to how much is already known) to form the updated predictions. These levels are coupled to each other by parameters ***κ*, *ω*** (between the first and second levels), and a constant ***ϑ*** determines the variance step-size for the third level. ***κ*** denotes a phasic, temporally varying parameter that determines scaling of the volatility, and ***ω*** and ***ϑ*** denotes a tonic, time-invariant parameter for the second and third level respectively which determines the sensitivity to update by each layer

The first level can be expressed as a Gaussian distribution whose mean is the previous value of the input and a variance which is a function of the level above.

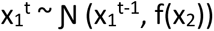

where the subscript denotes the level, and the superscript denotes the time. Similarly, for the ascending levels

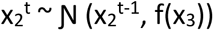

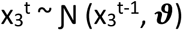

Note the variance of the final layer is determined solely from the parameter ***ϑ***.

The function that determines the coupling between the levels is denoted by

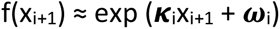

where ***κ*** is the phasic parameter that scales the variance and ***ω*** is the tonic, time-invariant parameter.

Having described the model hierarchies and their coupling, we can now describe how the posterior means of each of these levels (i.e., the beliefs) gets updated dynamically. The beliefs on each level, at every time t, are updated by

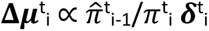

where Δ***μ***^t^_i_ is the update at time t for the posterior mean of the level i, which is dependent on the estimated precision (pi-hat) of the level below (new learning) and precision of current belief of the same level pi, and ***δ***^t^_i_ , the prediction error.

## Data/Code availability statement

The stimulus used and datasets generated during the analysis pipelines in the present study are available from the corresponding author on reasonable request.

## Acknowledgements

We acknowledge the generous support of NBRC Core funds and the Computing facility. This study was supported by Ramalingaswami Fellowship (Department of Biotechnology, Government of India) to DR (BT/RLF/Re-entry/07/2014) and grant number F.NO. K-15015/42/2018/SP-V from Ministry of Youth Affairs and Sports, Government of India to AB. DR was also supported by SR/CSRI/21/2016 extramural grant from the Department of Science and Technology (DST) Ministry of Science and Technology, Government of India. DR and AB acknowledge the generous support of the NBRC Flagship program BT/ MEDIII/ NBRC/ Flagship/ Program/ 2019: Comparative mapping of common mental disorders (CMD) over lifespan. Data collection and sharing for this project was provided by the Cambridge Centre for Ageing and Neuroscience (CamCAN). CamCAN funding was provided by the UK Biotechnology and Biological Sciences Research Council (grant number BB/H008217/1), together with support from the UK Medical Research Council and University of Cambridge, UK. In accordance with the data usage agreement for CamCAN dataset, the article has been submitted as open access.

## Ethics statement

CamCAN dataset was collected in compliance with the Helsinki Declaration, and has been approved by the local ethics committee, Cambridgeshire 2 Research Ethics Committee (reference: 10/H0308/50)

---

**Supplementary Figure 1:**
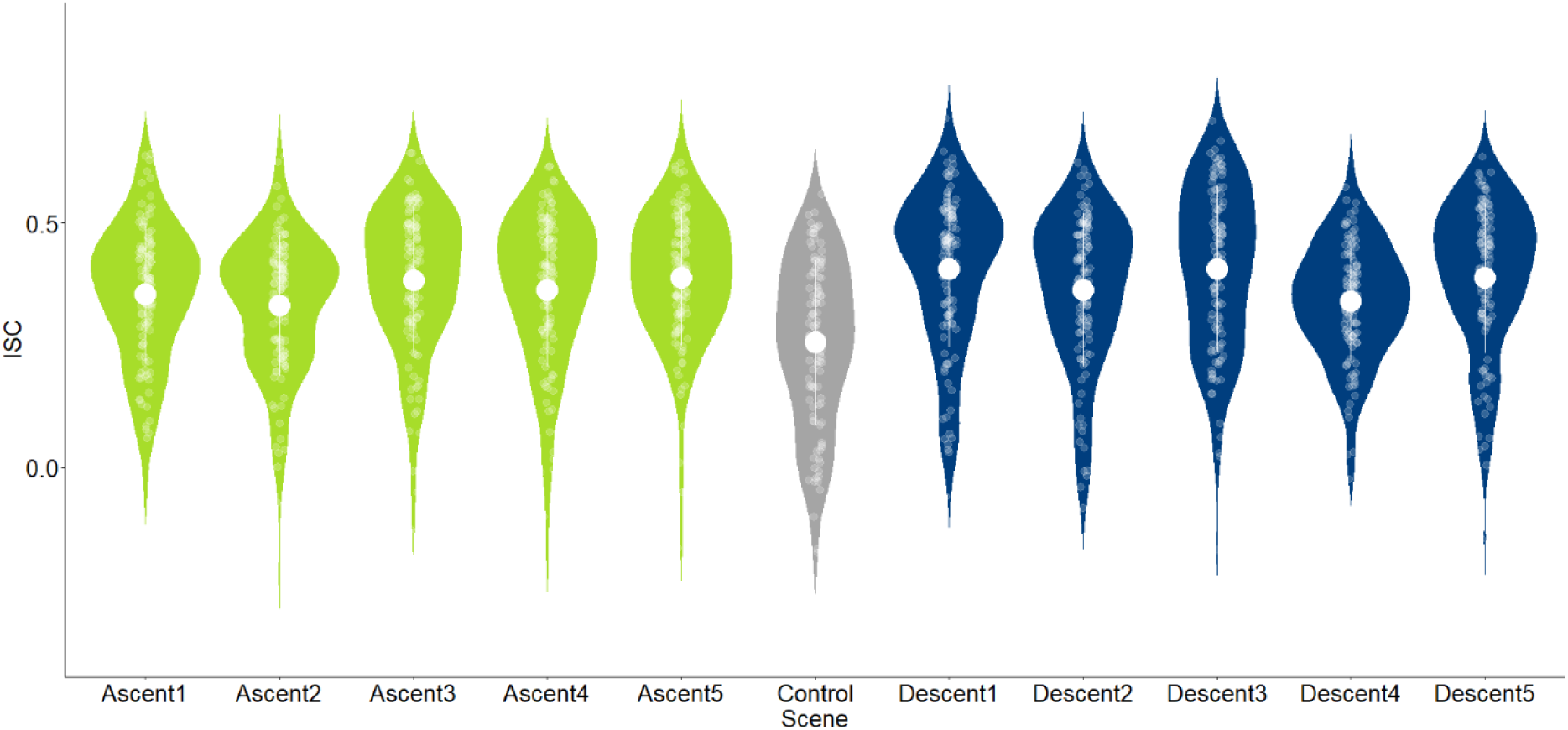
Scene-wise whole-brain ISC. All separate emotional scenes taken for analysis had significantly higher ISC than control segment. Ascent1>Control t = 4.7877, p-value = 5.294e-06, CI [0.058 0.140], Cohen’s d = 0.64, Ascent2>Control t = 3.6841, p-value = 0.0003577, CI [0.034 0.115], Cohen’s d = 0.48, Ascent3>Control t = 6.1302, p-value = 1.402e-08, CI [0.086 0.169], Cohen’s d = 0.80, Ascent4>Control t = 5.5561, p-value = 1.943e-07 CI [0.069 0.146], Cohen’s d = 0.66, Ascent5>Control t = 7.0599, p-value = 1.58e-10, CI [0.095 0.170], Cohen’s d = 0.86. Descent1>Control t = 7.5092, p-value = 1.67e-11 CI [0.110 0.188], Cohen’s d = 0.91, Descent2>Control t = 4.9096, p-value = 3.192e-06 CI [0.064 0.150], Cohen’s d= 0.66, Descent3>Control t = 7.1994, p-value = 7.902e-11 CI [0.108 0.191], Cohen’s d = 0.89, Descent4>Control t = 4.4168, p-value = 2.355e-05 CI [0.046 0.121], Cohen’s d = 0.57, Descent5>Control t = 6.1723, p-value = 1.151e-08, CI [0.089 0.174], Cohen’s d =0.82

**Supplementary Figure 2:**
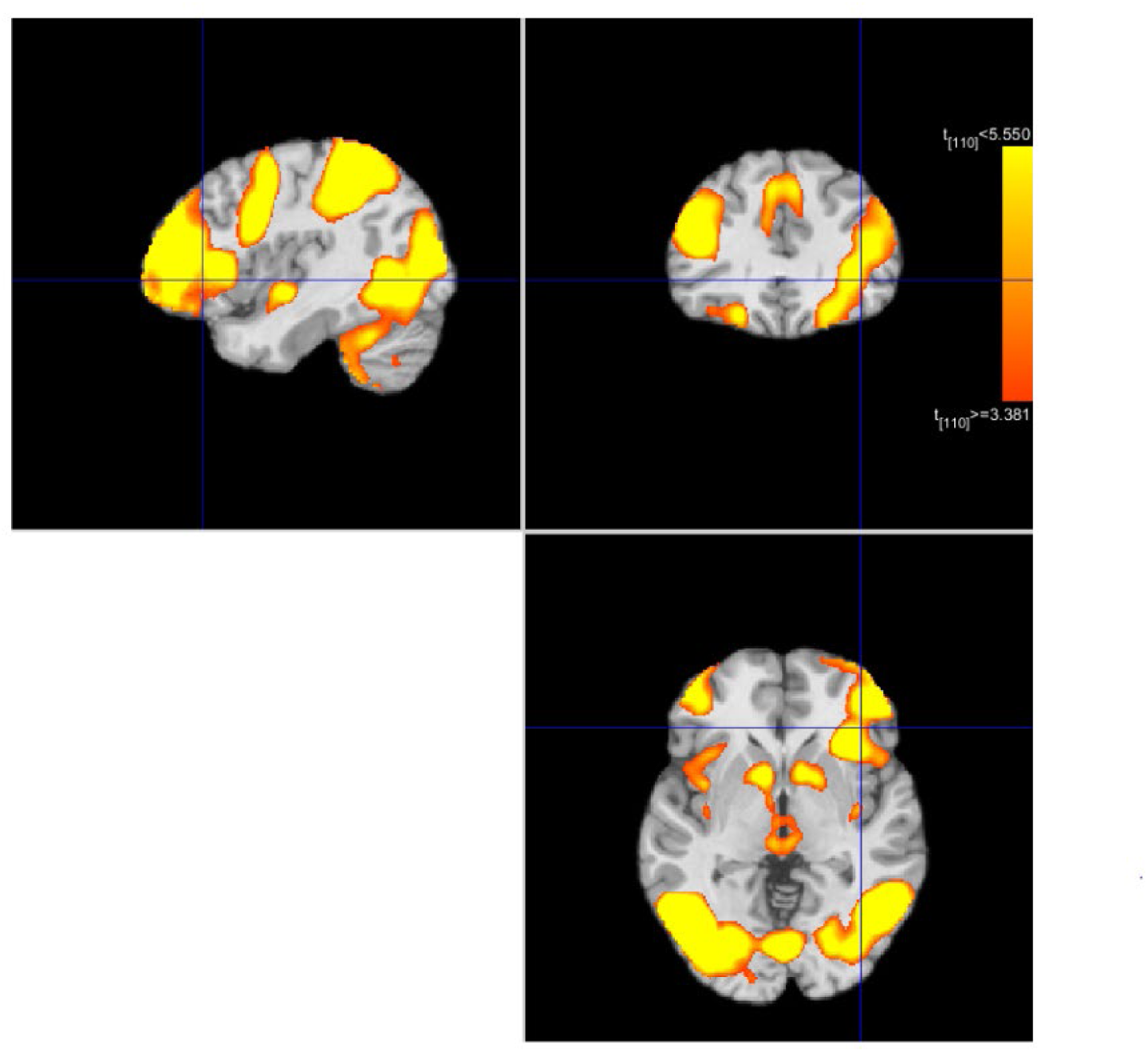
Whole brain maps of Ascent-3 showing clusters of neural activation (n=111) for Ascent>Control. This contrast map is a representative contrast map showing the ROI localisations for one Ascent. We selected significant clusters of p < 0.001 with voxel-extent threshold of k > 25.

**Supplementary Figure 3:**
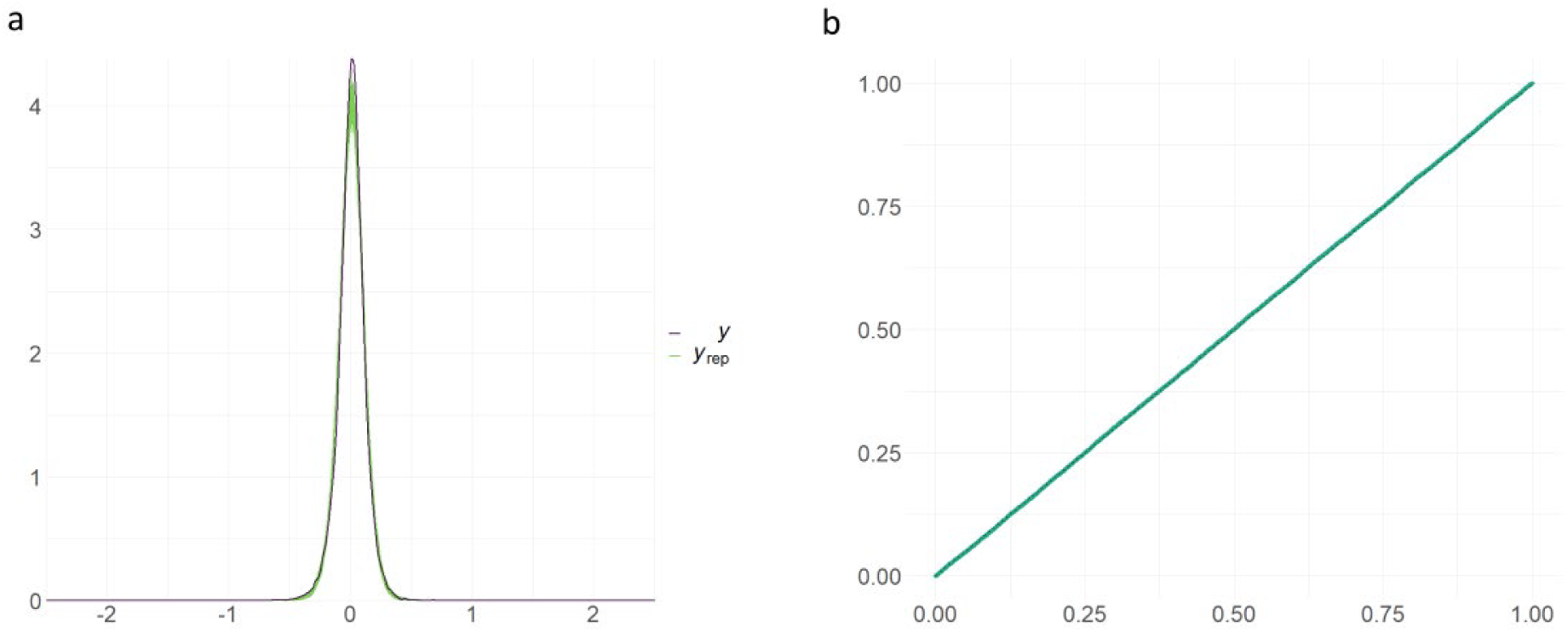
Model fits. (**a**) Posterior predictive check (PPC) for the main model in analysis suggesting overall good model fit. (data in black, simulated samples in green). (**b**) To check for robustness of model to outliers, model calibration using cross validation was performed by comparing the probability integral transform (PIT) checks to a standard uniform distribution (dashed line). Both results were obtained using 1000 draws from the model.

**Supplementary Figure 4:**
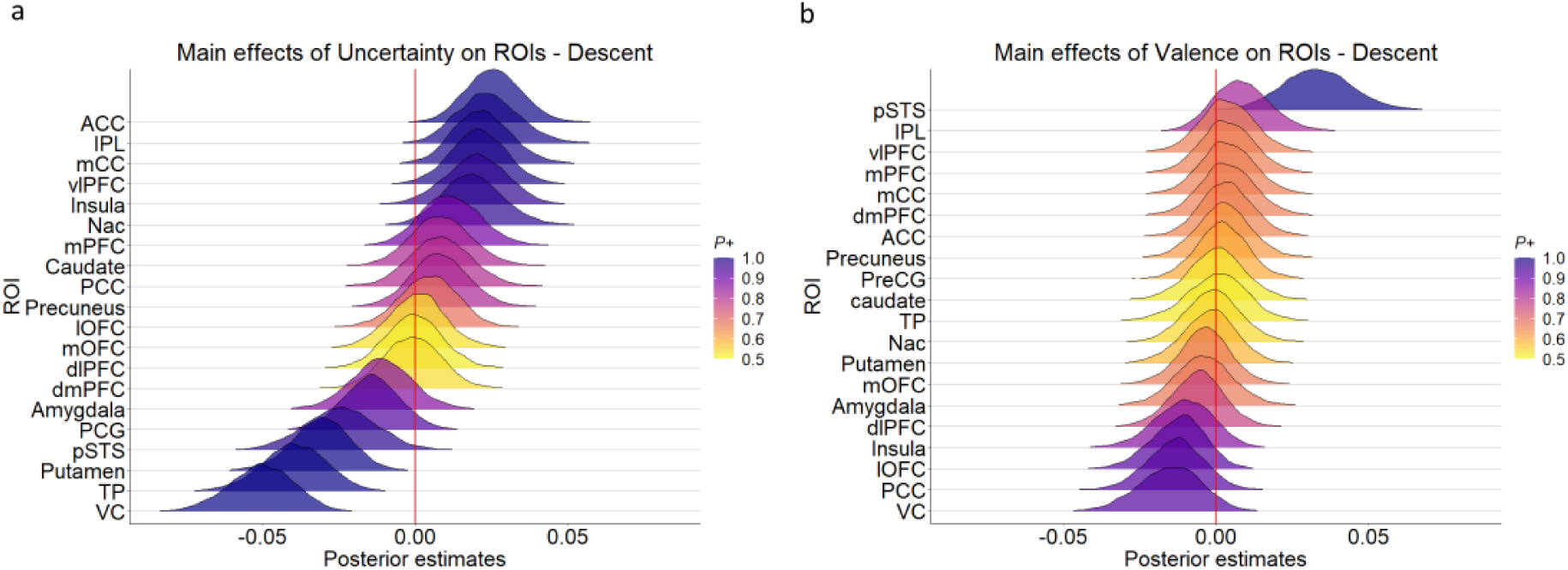
Bayesian hierarchical regression (Descent). More regions were modulated by Uncertainty (**a**) in Descent (as was observed in Ascent), compared to Valence (**b**).

**Supplementary Figure 5:**
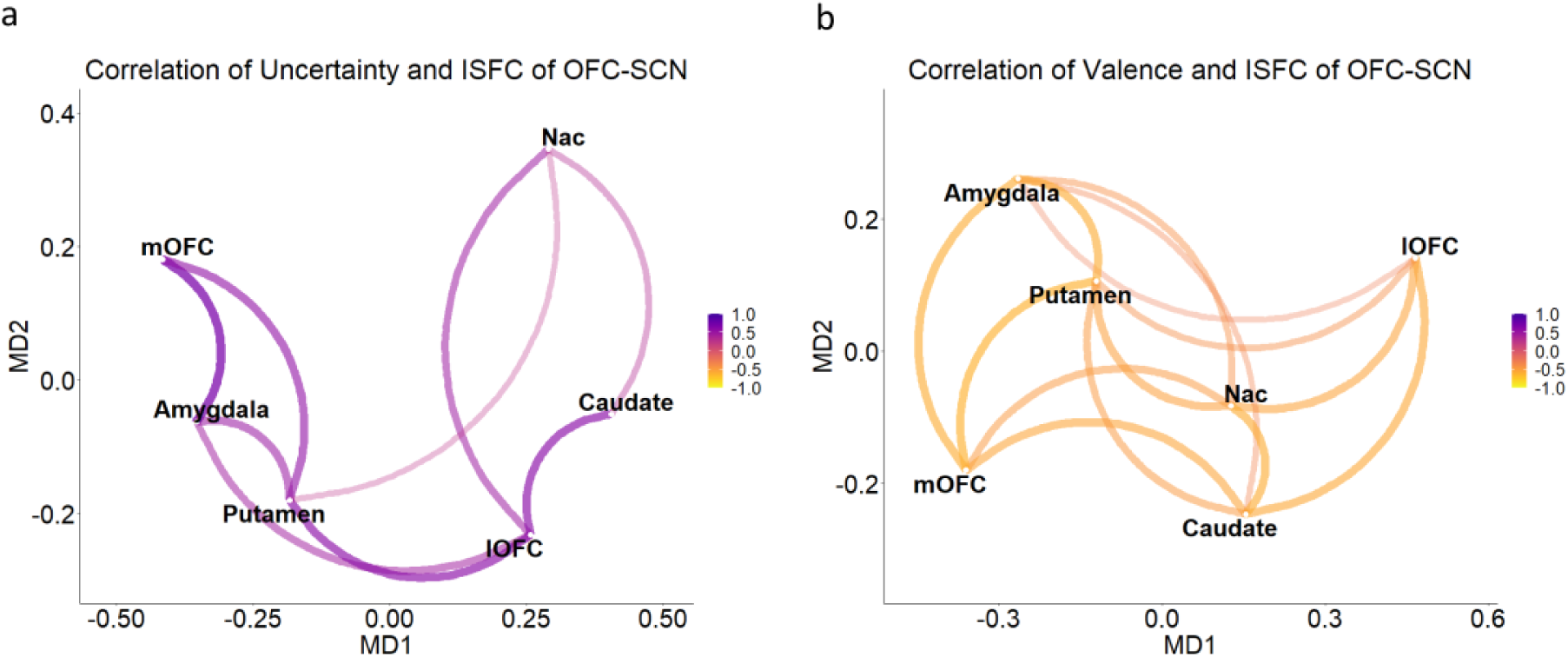
Multidimensional scaling of ISFC and Uncertainty correlation. ISFC of OFC and subcortical network (Amygdala, Nucleus Accumbens, Caudate and Putamen) correlated with Uncertainty (**a**) and Valence (**b**).

**Supplementary Figure 6:**
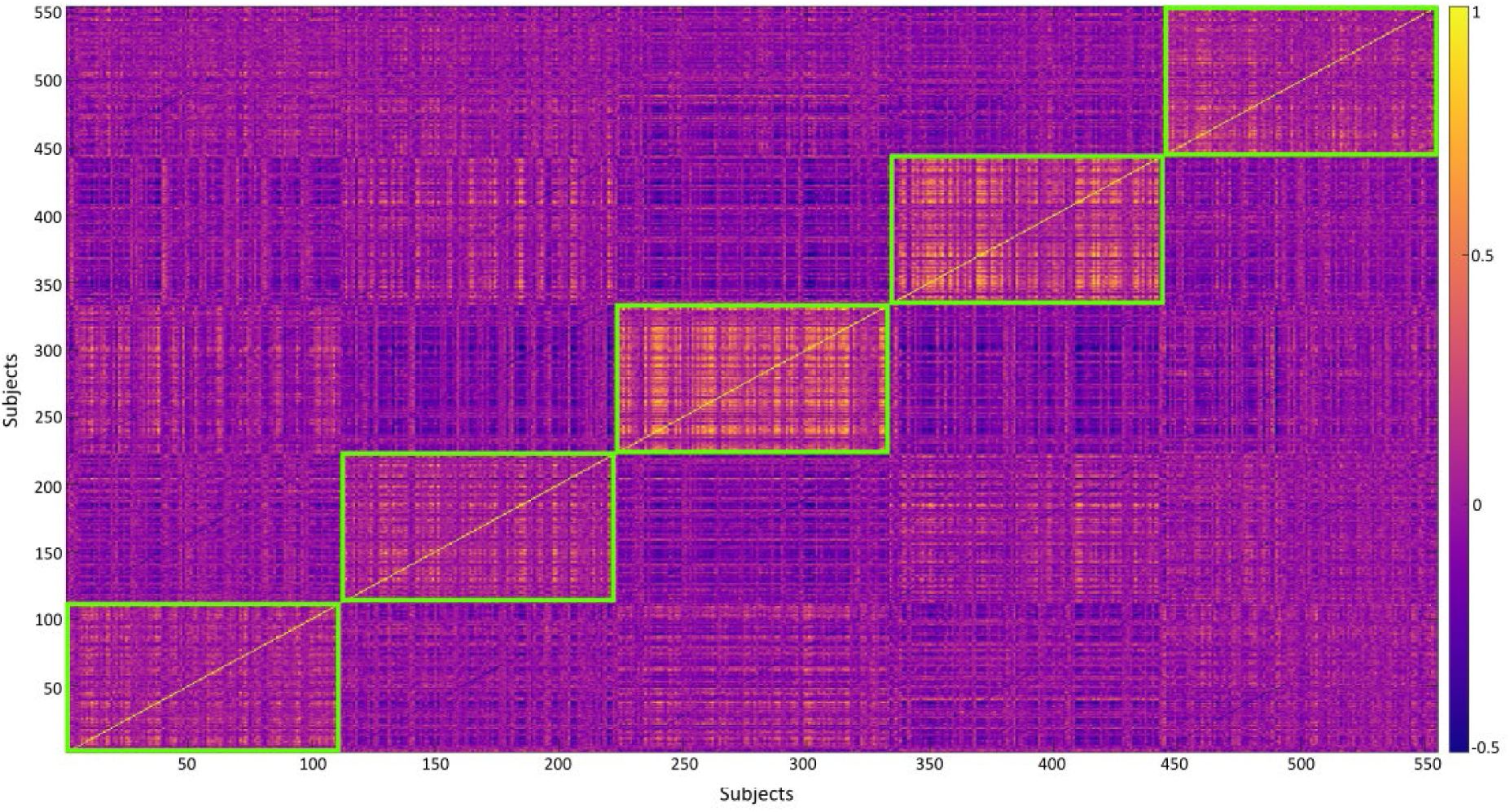
Individual HMM. Single subject HMM conducted for lOFC shows consistency of states across subjects. State 1 mean r = 0.27, State 2 mean r = 0.29, State 3 mean r = 0.45, State 4 mean r = 0.40, State 5 mean r = 0.29.

**Supplementary Figure 7:**
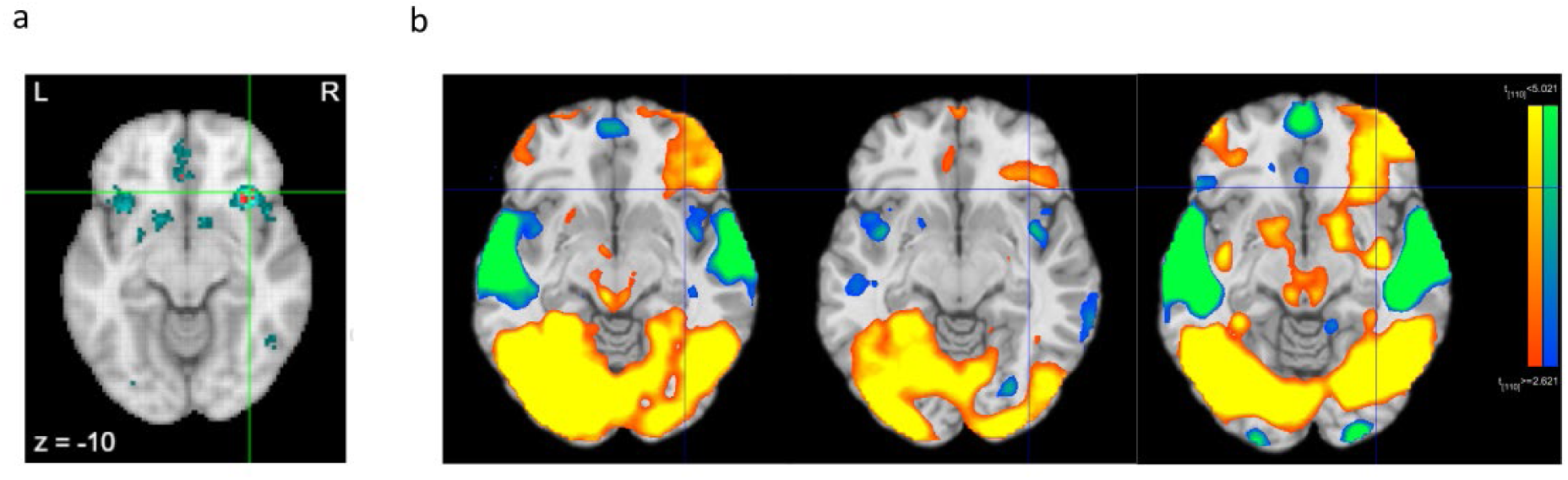
(**a**) Neurosynth coordinates for term-based search of ‘uncertainty’ with uniformity maps and association maps (formerly forward and reverse inference maps). Uniformity maps (blue) show voxels consistently activated in studies using the term uncertainty (130 studies), and association maps (red) show areas specific to such studies compared to studies without the term. (**b**) The same coordinate in lOFC in three Ascents in our study.

**Supplementary Table 1:** Hierarchical Gaussian Filter – Prior parameters for the 4 models. *µ*_n_ mean of layer n, *σ*_n_ variance of layer n(logspace) (both layer mean and variance have hyperpriors on them with mean *µ* and variance *σ*), *ρ*_n_ *µ* mean of drift parameter of layer n, *ρ*_n_ *σ* variance of drift parameter of layer n, *κ*_n_ *µ* mean of phasic parameter (logspace), *κ*_n_ *σ* variance of phasic parameter (logspace), *ω*_n_ *µ* mean of tonic parameter of layer n, *ω*_n_ *σ* variance of tonic parameter of layer n, *θ µ* mean of volatility parameter of third layer, *θ σ* variance of volatility parameter of third layer, *π µ* mean of precision (logspace), *π σ* variance of precision (logspace).

| Parameters | M1 2-HGF Low | M2 2-HGF High | M3 3-HGF Low | M4 3-HGF High |
| --- | --- | --- | --- | --- |
| $\mu_1 \mu$ | 99991 | 99991 | 99991 | 99991 |
| $\mu_2 \mu$ | 1 | 1 | 1 | 1 |
| $\mu_3 \mu$ | - | - | 0 | 0 |
| $\mu_1 \sigma$ | 99992 | 99992 | 99992 | 99992 |
| $\mu_2 \sigma$ | 0 | 0 | 0 | 0 |
| $\mu_3 \sigma$ | - | - | 0 | 0 |
| $\sigma_1 \mu$ | 99993 | 99993 | 99993 | 99993 |
| $\sigma_2 \mu$ | log(0.1) | log(0.1) | log(0.1) | log(0.1) |
| $\sigma_3 \mu$ | - | - | log(0.1) | log(0.1) |
| $\sigma_1 \sigma$ | 0 | 0 | 0 | 0 |
| $\sigma_2 \sigma$ | 0 | 0 | 0 | 0 |
| $\sigma_3 \sigma$ | - | - | 0 | 0 |
| $\rho_1 \mu$ | 0 | 0 | 0 | 0 |
| $\rho_2 \mu$ | 0 | 0 | 0 | 0 |
| $\rho_3 \mu$ | - | - | 0 | 0 |
| $\rho_1 \sigma$ | 0 | 0 | 0 | 0 |
| $\rho_2 \sigma$ | 0 | 0 | 0 | 0 |
| $\rho_3 \sigma$ | - | - | 0 | 0 |
| $\kappa_1 \mu$ | log(0.2) | log(2) | log(.2) | log(2) |
| $\kappa_2 \mu$ | - | - | log(.01) | log(.1) |
| $\kappa_1 \sigma$ | 0 | 0 | 0 | 0 |
| $\kappa_2 \sigma$ | - | - | 0 | 0 |
| $\omega_1 \mu$ | 99993 | 99993 | 99993 | 99993 |
| $\omega_2 \mu$ | -8 | -8 | -8 | -8 |
| $\omega_1 \sigma$ | 1 | 1 | 1 | 1 |
| $\omega_2 \sigma$ | .05 | .05 | .05 | .05 |
| $\vartheta \mu$ | - | - | 2 | 2 |
| $\vartheta \sigma$ | - | - | .05 | .05 |
| $\pi \mu$ | -99993 | -99993 | -99993 | -99993 |
| $\pi \sigma$ | 4 | 4 | 4 | 4 |

**Supplementary Table 2:** Hierarchical Gaussian Filter – Posterior parameters for the 4 models

| Parameters | M1 2-HGF Low | M2 2-HGF High | M3 3-HGF Low | M4 3-HGF High* |
| --- | --- | --- | --- | --- |
| $\mu_1$ | -0.037 | -0.037 | -0.037 | -0.033 |
| $\mu_2$ | 1 | 1 | 1 | 1 |
| $\mu_3$ | - | - | 0 | 0 |
| $\sigma_1$ | 0.00036 | 0.00036 | 0.00036 | 0.00036 |
| $\sigma_2$ | 0.1 | 0.1 | 0.1 | 0.1 |
| $\sigma_3$ | - | - | 0.1 | 0.1 |
| $\rho_1$ | 0 | 0 | 0 | 0 |
| $\rho_2$ | 0 | 0 | 0 | 0 |
| $\rho_3$ | - | - | 0 | 0 |
| $\kappa_1$ | 0.2 | 2 | 0.2 | 2 |
| $\kappa_2$ | - | - | .01 | .1 |
| $\omega_1$ | -8.886 | -10.458 | -8.889 | -10.890 |
| $\omega_2$ | -7.778 | -4.331 | -7.776 | -6.888 |
| $\vartheta$ | - | - | 2 | 3.37 |
| $\pi$ | 2085088 | 34344044 | 2091958 | 12456708 |
| Log model evidence (LME) | 1370 | 1520 | 1370 | 1577 |
| Correlation with Uncertainty | -0.14 | 0.11 | -0.25 | 0.46 |

**Supplementary Table 3.**
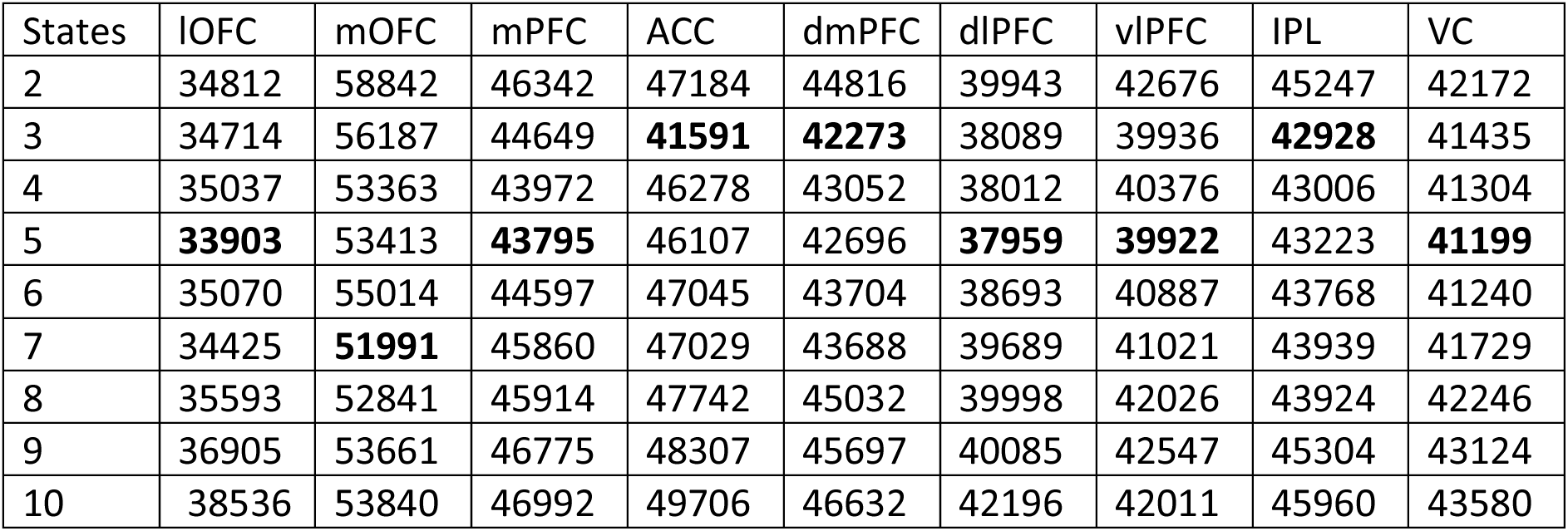
Bayesian Information Criteria (BIC) scores for states [2, 10] for different ROIs. Chosen number of states correspond to minimal BIC scores for all ROIs taken.

